# *Staphylococcus Aureus* Colonization and Familial Transmission Over a One Year Period

**DOI:** 10.1101/144782

**Authors:** Blake M. Hanson, Ashley E. Kates, Sean M. O’Malley, Elizabeth Mills, Loreen A. Herwaldt, James C. Torner, Jeffrey D. Dawson, Sarah A. Farina, Cassandra Klostermann, Tara C. Smith

## Abstract

**Methods:** 177 adults and 86 minors comprising 95 family units were enrolled from two counties in Iowa and followed up for 52 weeks. Random effects logistic regression was used to test the effect of different risk factors on the probability of an individual falling into a different *S. aureus* colonization categories. Additionally, the frequency of *S. aureus* colonization events and familial transmission events were calculated.

**Results:** The number of positive environmental sites within a participant’s house was associated with being a persistent carrier compared to being a non-carrier or intermittent carrier. Age, sharing bath towels, and the number of positive environmental sites within a participant’s house were associated with being a persistent or intermittent carrier. Colonization events per year were 3.95 for adults and 3.04 for minors. Duration of colonization was longest for persistent carriers (92.3 days for adults and 97.8 days for minors), and intermittent carriers had the most colonization events.

**Conclusions:** The average duration of colonization was significantly different when comparing intermittent carriers and non-carriers. We have also established estimates of the duration of colonization and the frequency of transmission events among family units in a non-healthcare population.

## Introduction

*Staphylococcus aureus* (*S. aureus*) is a commensal bacterium, with the nares historically considered the most common colonization site (1, 2). About 30% of the population is colonized with *S. aureus* (3), and most colonized persons are asymptomatic carriers. Asymptomatic carriage may not harm the host, but is a known risk for subsequent symptomatic infections in both the healthcare and community settings (4).

Numerous prior studies have assessed *S. aureus* colonization of the nares and oropharynx in the healthcare setting (5–9), while few have assessed colonization of these sites in a population of healthy community members (10–12). Moreover, we did not identify any published studies that estimated the frequency of colonization events or familial transmission of *S. aureus* in the absence of an infected family member.

This study assessed data on *S. aureus* colonization collected from a longitudinal cohort of 95 family units that was recruited from rural and urban Iowa and had minimal healthcare contact. Utilizing data from this cohort, we determined the incidence of colonization with *S. aureus*, the duration of colonization, the frequency of familial transmission, and the incidence of infections. We also, characterized the colonizing *S. aureus* strains, and we assessed risk factors for colonization.

## Methods

### Study design and sample collection

Participants were enrolled in a prospective cohort study. Ninety-five families from two counties in Iowa, Johnson County and Keokuk County, were enrolled between October 6^th^, 2011 and January 4^th^ 2012. These two counties were chosen to reflect an urban (Johnson County) and a rural population (Keokuk County) in Iowa, as defined by the US Census Bureau (13).

County-specific methodologies were employed to recruit potential participants. Residents in Johnson County were recruited via advertisements in a local newspaper and by mailing lists. Interested persons contacted our study team via email. Participants from Keokuk County were recruited from the Keokuk Rural Health Study, a previously existing cohort (14), and were contacted via letter; the enrollment study visit was scheduled via subsequent phone call.

Trained study team members traveled to each family’s home to enroll the family members. During the study visit, potential participants were screened to assess enrollment criteria, instructed on sample collection techniques, and completed enrollment interviews. Inclusion criteria for enrollment in the cohort were: participant must be able to provide consent, assent, or have parents willing to provide consent; and must be willing to participate by completing enrollment and weekly follow-up questionnaires. All participants signed an informed consent document or assent document prior to completing any study activities. The University of Iowa institutional review board approved all study protocols prior to recruitment.

Participants completed questionnaires collecting data on demographics, healthcare exposure, farming exposure, medical conditions, and other risk factors for *S. aureus* colonization and infection. When collecting weekly samples, participants also completed a questionnaire about potential skin and soft tissue infections.

### Isolation of *S. aureus*

While at each family’s home, study team members instructed each enrolled family member on the proper technique for collecting nasal samples. Adult participants were instructed on providing oropharyngeal samples as well. Participants were also given BBL CultureSwabs so they could sample possible *S. aureus* infections. Samples were collected with BBL CultureSwabs that included Liquid Stuart Medium (Becton, Dickinson and Company, Sparks MD, USA). Samples were transported through the US Postal system with icepacks to the Center for Emerging Infectious Diseases (CEID) laboratory facilities.

Study team members collected environmental swabs during the enrollment visit using 3-inch by 4-inch sterile duster cloths. Six commonly touched surfaces (living room television remote, main bathroom toilet flush lever, main bathroom light switch, kitchen sink handle, refrigerator door handle, and oven knobs) were sampled by wiping the surfaces in all directions for one minute. Each site was sampled with a new duster cloth, and the cloth was placed into a dry sterile bag. The samples were transported to the CEID on ice.

Nasal and oropharyngeal swabs were inoculated separately into 5mL of Baird-Parker medium and incubated at 35°C for 24 hours. Environmental samples were processed by adding 25mL of 1.0% peptone to the bag and homogenizing by hand for one minute. 5mL of the 1.0% peptone was added to 5mL of 2x Baird-Parker medium and incubated at 35°C for 24 hours, Following incubation, isolates were plated onto Baird-Parker agar and BBL CHROMagar MRSA II (Becton, Dickinson and Company, La Jolla CA, USA) and incubated for 48 hours at 35°C. Presumptive *S. aureus* colonies (mauve colonies on CHROMagar MRSA II plates, and black colonies with shiny halos on Baird-Parker agar) were streaked onto Columbia CNA agar with 5% sheep’s blood (Becton, Dickinson and Company, Sparks MD, USA) and incubated for 24 hours at 35°C.

### Characterization of *S. aureus*

Presumptive *S. aureus* colonies grown on Columbia CNA agar with 5% sheep’s blood were assessed with the catalase test, coagulase test, and Pastorex Staph Plus rapid latex agglutination assay (Bio-Rad, Redmond WA, USA). All isolates confirmed to be *S. aureus* had genomic DNA extracted for molecular testing.

### Antimicrobial susceptibility testing

Phenotypic resistance to antimicrobials was assessed through minimum inhibitory concentration (MIC) testing following the Clinical and Laboratory Standards Institute’s standards (15). Isolates were tested for susceptibility to oxacillin, tetracycline, erythromycin, clindamycin, trimethoprim-sulfamethoxazole, gentamycin, levofloxacin, vancomycin, daptomycin, quinupristin/dalfopristin, linezolid, and rifampin.

### Molecular testing

Bacterial genomic DNA was extracted with the Promega Wizard Genomic DNA purification kit (Promega Corporation, Madison WI, USA). Polymerase chain reaction (PCR) was used to assess the presence of the *mec*A gene (16), and the genes encoding Panton-Valentine leukocidin (PVL) (17), and to amplify the protein A (*spa*) gene (18). Strain type was identified through typing the *spa* gene using the Ridom StaphType software (Ridom GmbH, Germany). Isolates that could not be typed three times were labeled as non-typable (NT) (19). The Based Upon Repeat Pattern (BURP) algorithm was used to identify and characterize genetic clusters (20). All molecular procedures included known positive and negative controls.

### Statistical analysis

Data were analyzed using SAS software version 9.3, and R version 2.15.3. Demographic data were assessed for differences between the two counties using the Student’s T-Test to assess age, Chi-Squared Test without Yate’s Continuity Correction to assess gender, and Fisher’s Exact Test to assess dichotomous variables with small sample sizes.

Participant carrier status was determined by taking the number of nares and oropharyngeal swabs that grew *S. aureus* divided by the total number of swabs over the study period, excluding the baseline cultures and culture results. The cutoffs utilized in this manuscript were previously described by van Belkum et al., but were modified to include the oropharynx in the definition (21). Participants were classified as: ‘non-carriers’ if ≤10% of samples from both the nares and the oropharynx grew *S. aureus*, ‘intermittent carriers’ if between 10% and 80% of samples from either the nares or the oropharynx grew this organism, and ‘persistent carriers’ if ≥80% of samples from either the nares or the oropharynx grew *S. aureus*.

Logistic regression with receiver operating curves (ROC) was used to determine the lowest number of nares and oropharynx swabs necessary to accurately predict each participant’s carrier status. For adults, the number of culture positive nares swabs and the number of culture positive oropharynx swabs were modeled as separate variables. The ROC analysis was completed separately for adults and minors, as minors submitted nares swabs only, to generate true positive rates (TPR) and false positive rates (FPR).

Random effects logistic regression using SAS proc glimmix with Satterthwaite’s approximation for degrees of freedom was used to test the effect of risk factors on: the probability of being an intermittent carrier given one is not a persistent carrier, the probability of being a persistent carrier compared with non-carriers and intermittent carriers, and the probability of being an intermittent carrier or persistent carrier compared with non-carriers. Number of children, age, and house size were modeled as continuous variables. All other risk factors were modeled as dichotomous or continuous variables. Risk models for adults were adjusted for age; models for minors were adjusted for age, gender, and an age and gender interaction term. Random effects logistic regression was used to assess the random effects of both family units and county in addition to the effects of specific risk factors. Some models had a non-positive variance for county after adjustment for the random effect of family and the risk factor, indicating that the remaining variance was not attributable to county, and thus the variance was set to zero in these instances (22).

Colonization events and familial transmission events were identified based upon the following definitions. A colonization event was defined as a positive nares or oropharyngeal culture (i.e., acquisition of carriage), when the participant’s samples from the week prior were negative in both the nares and the oropharynx. For each colonization event, the duration of colonization was calculated as the time from the beginning of the colonization event (i.e., the first positive culture after a prior negative set of nares and oropharyngeal cultures) until the time cultures of both the nares and oropharynx swabs were negative concurrently. If a participant was colonized during the first week of observation, or during the final week of observation, left truncation and right truncation of the interval were used, respectively. All other colonization durations were calculated with interval censoring. For example, if a participant’s nares and the oropharyngeal cultures were negative during one sampling, and were positive at the next two samplings seven days apart, and were negative seven days after that, half of the first interval and half of the last interval (3.5 days each) would be added to the 14 days between the two sets of positive cultures for a total colonization duration of 21 days. If a participant began the study with a positive culture, had a positive culture seven days later, and had two negative cultures seven days after that, the colonization duration would be 17.5 days. The patient would accrue seven days for the positive culture at enrollment, seven days for the second positive culture, and 3.5 days for the right truncation. The duration would be left truncated because we do not know how many weeks prior to our study the participant had positive cultures. We used these censoring and truncation methodologies estimate the duration of colonization most accurately without overestimating this time period. Familial transmission was defined as the acquisition of colonization with a specific strain of *S. aureus* found concurrently within a family member. Concurrent colonization was defined as a current culture from a family member that grew a specific strain of *S. aureus* that was also currently colonizing a different family member, indicating two family members were colonized by the same strain at the same time. The familial transmission event occurs when the second family member acquires the strain.

## Results

The logistic regression models with ROC curves indicated the minimum number of swabs necessary to maximize the TPR while minimizing the FPR for adults was 14 consecutive sets of swabs for adults, and a single swab for minors. Of the 177 adults, 154 (87.0%) submitted 14 or more swabs after the baseline swab and were included in this analysis. Of the 86 minors, 77 (89.5%) continued past the baseline and were included in analysis. Adults submitted an average of 34.71 swabs (range: 14-52) and minors submitted an average of 30.87 swabs (range: 2-46). The average age of the adult participants was 44.5 (23.10-67.72 years), while the average age of minors was 9.65 years (0.42-18.18 years). Additional demographics information is included in the supplemental Tables C2 and C3.

A total of 10,987 swabs were processed during follow-up, 3,119 of which were culture positive for *S. aureus*. Among adults, 29.8% of cultures from at least one anatomical site grew methicillin-susceptible *S. aureus* (MSSA), and 1.3% of cultures grew MRSA. Among minors, 30.4% of nares cultures grew MSSA, and 0.6% of nares cultures grew MRSA. For additional details on prevalence, see Table 1. Of the adults, 47.4% met the definition of non-carrier, 32.2% met the definition of intermittent carrier, and 20.4% met the definition of persistent carrier (Table 3, Figure 1). Additional details on site-specific prevalence can be found in Table S1.

**Figure 1:**
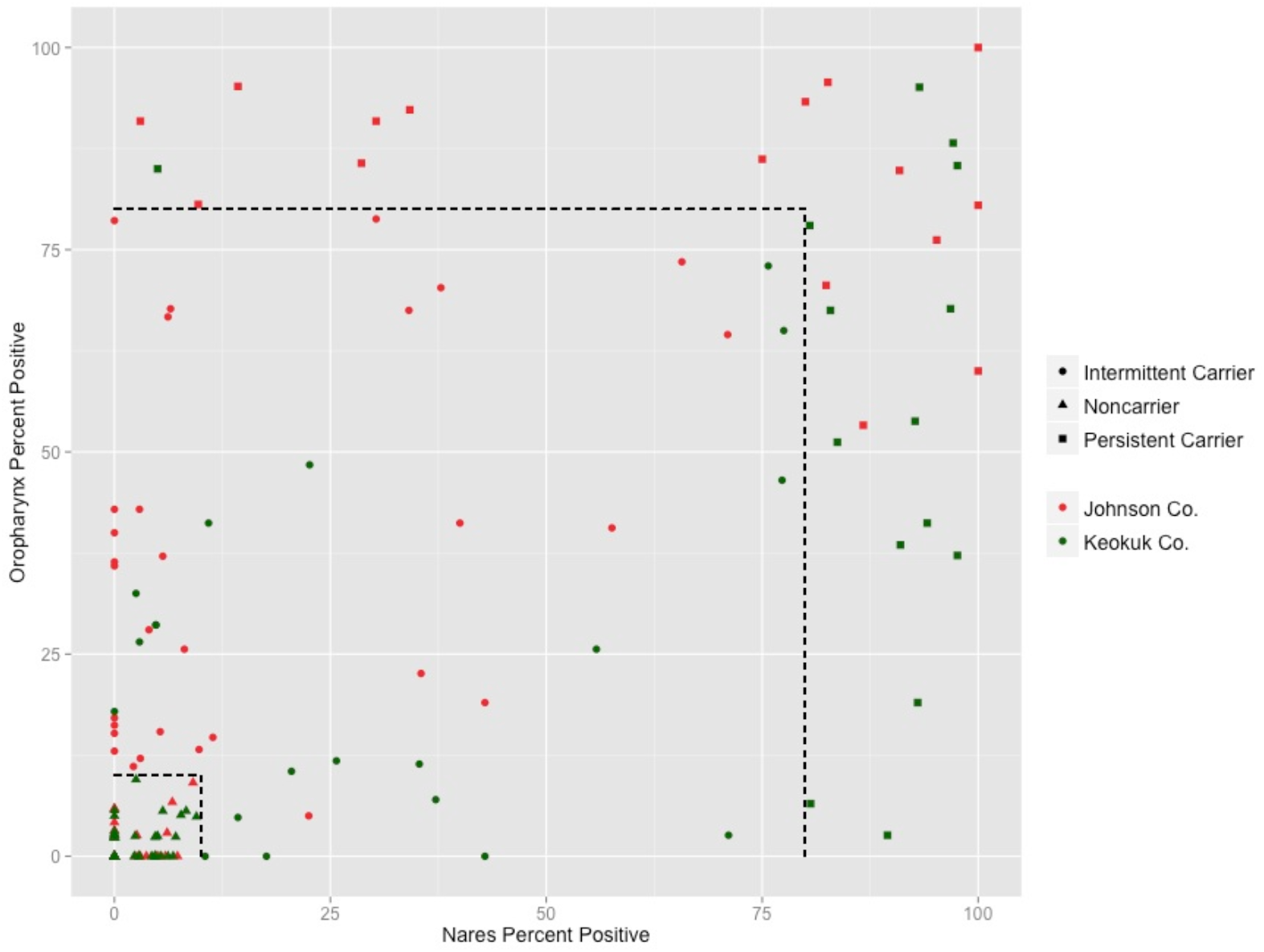
Scatterplot of participant nares and oropharynx colonization rates. Each circle denotes a single participant. Black-hashed lines indicate cutoffs between non-carriers, intermittent carriers, and persistent carriers (21).

**Table 1:** Total number of sets of oropharyngeal and nasal cultures that grew MSSA or MRSA, age grouped by county

|  | MSSA | MRSA<sup>*</sup> | Total |
| --- | --- | --- | --- |
| **Adults** |  |  |  |
| **Johnson Co.** | 820/2228 (36.8%) | 12/2228 (0.5%) | 832/2228 (37.3%) |
| **Keokuk Co.** | 849/2904 (29.2%) | 55/2904 (1.9%) | 904/2904 (31.1%) |
| **Total** | 1669/5132 (32.5%) | 67/5132 (1.3%) | 1736/5132 (33.8%) |
| **Minors** |  |  |  |
| **Johnson Co.** | 405/1292 (31.3%) | 12/1292 (0.9%) | 417/1292 (32.3%) |
| **Keokuk Co.** | 294/1052 (27.9%) | 2/1052 (0.2%) | 296/1052 (28.1%) |
| **Total** | 699/2344 (29.8%) | 14/2344 (0.6%) | 713/2344 (30.4%) |

Note: The numerator for adults was the number of participants with *S. aureus* isolated from the nares, the oropharynx, or both. The numerator for minors was the number of participants with *S. aureus* isolated from the nares. The denominator is the total number of participants by county for adults and minors.
<sup>\*</sup>MRSA isolates are designated by the presence of the *mecA* gene.

One hundred twenty-five *spa* types were identified amongst the 3,119 *S. aureus* isolates, of which, the most common were t002 (n=345), t008 (n=248), and t216 (n=180). Adjustment for family clonal isolates was done by taking a single representative sample of each clonal strain from each family unit. Following adjustment, the number of unique isolates was reduced to 278. The most common *spa* types following adjustment were t002 (n=23), t084 (n=17), t008 (n=16), and t216 (n=16). The BURP algorithm identified 10 clusters in Keokuk County and seven clusters in Johnson County, but the number of *spa* types per cluster was higher in Johnson County than in Keokuk County (Appendix Figures S1 and S2). After adjustment, *mec*A prevalence was 5.4%, and PVL prevalence was 1.8% (Table 2).

**Table 2:** Molecular characteristics of *S. aureus* isolates

|  | *mecA* |  | PVL |  |
| --- | --- | --- | --- | --- |
|  | Adjusted | Unadjusted | Adjusted | Unadjusted |
| **Johnson Co.** | 5/156 (3.2%) | 24/1569 (1.5%) | 0/156 (0%) | 0/1569 (0%) |
| **Keokuk Co.** | 10/122 (8.2%) | 57/1550 (3.7%) | 5/122 (4.1%) | 26/1550 (1.7%) |
| **Total** | 15/278 (5.4%) | 81/3119 (2.6%) | 5/278 (1.8%) | 26/3119 (0.8%) |

Note: Unadjusted values include all *S. aureus* isolates from participants broken down by county. Adjusted values include only a single copy of isolates with the same molecular profile (same *spa* type, and presence/absence of *mecA* and PVL) to account for familial clustering and the presence of clonal isolates in the nares and oropharynx of adult participants.

**Table 3:** Participants by carrier group

|  | **Johnson Co.**<br>**(n=73)** | **Keokuk Co.**<br>**(n=79)** | **Total**<br>**(n=152)** |
| --- | --- | --- | --- |
| **Non-carrier** | 28 (38.36%) | 44 (55.70%) | 72 (47.37%) |
| **Intermittent carrier** | 29 (39.73%) | 20 (25.32%) | 49 (32.24%) |
| **Persistent carrier** | 16 (21.92%) | 15 (18.99%) | 31 (20.39%) |

Note: The cutoffs used are as follows: non-carrier - $\leq 10\%$ of cultures from both the nares and the oropharynx were positive for *S. aureus*. Intermittent carrier - between 10% and 80% of cultures were positive from both the nares and the oropharynx. Persistent carrier - $\geq 80\%$ of cultures were positive from either the nares or the oropharynx.

Antimicrobial susceptibility testing was performed on a subset of 725 isolates obtained from samples received during the first week of each study month. Observed rates of resistance were zero for most tested antibiotics. However, 25.9% of isolates were resistant to erythromycin after adjustment and 17.7% were intermediately resistant to quinupristin/dalfopristin. Resistance rates to other agents were all under 10% (Figure 3).

Random effects logistic regression was used to assess the probability of being an intermittent carrier given that one was not a persistent carrier, and found significant associations with age (p-value: 0.0145) and with the number of positive environmental sites (p-value: 0.0137). Farming exposure did not quite meet the 0.05 significance level (p-value of 0.0733). The number of positive environmental sites was the only factor associated with being a persistent carrier compared with being a non-carrier or an intermittent carrier (p-value: 0.0137). Age (p-value: 0.0353), sharing bath towels (p-value: 0.0338), and the number of positive environmental sites (p-value: 0.0026) were associated with being either a persistent carrier or an intermittent carrier, compared with being a non-carrier. Data on the directionality of associations are presented in Tables 5 and 6.

The average number of observed colonization events was similar for adults (3.95) and minors (3.04). Among both adults and minors, intermittent carriers had the greatest number of observed colonization events. Duration of colonization was longest among persistent carriers: 92.3 and 97.8 days for adults and minors, respectively. Both adults and minors acquired strains carried by a family member (familial transmission), with an average of 0.77 events per person year of follow-up for adults, and 1.12 events for minors (Table 4). Participants reported skin and soft tissue infections during study follow-up, with six participants (four adults and two minors) reporting eight potential infections. Five wound swabs were submitted for culture, one (20%) of which grew *S. aureus*. The strain isolated from the wound and the participant’s current colonizing strain had the same spa type. The observed rate of putative infections per person year of follow-up was 0.049 for adults and 0.060 for minors.

**Table 4:** Colonization events and familial transmission events for adults and minors

|  | **Number of**<br>**Colonization**<br>**Events per**<br>**Person Year**<br>**Follow-up** | **Average**<br>**Colonization**<br>**Duration**<br>**(days)** | **Familial**<br>**Transmission**<br>**Events per**<br>**Person Year**<br>**Follow-up** |
| --- | --- | --- | --- |
| **Adults** |  |  |  |
| Non-carrier | 1.56 | 10.77 | 0.33 |
| Intermittent Carrier | 7.72 | 20.69 | 1.31 |
| Persistent Carrier | 3.64 | 92.30 | 0.92 |
| All Adults | 3.95 | 31.71 | 0.77 |
| **Minors** |  |  |  |
| Non-carrier | 1.24 | 7.07 | 0.62 |
| Intermittent Carrier | 5.39 | 25.71 | 1.89 |
| Persistent Carrier | 3.37 | 97.79 | 1.57 |
| All Minors | 3.04 | 35.31 | 1.22 |

Note: The cutoffs used are as follows: non-carrier - $\leq 10\%$ of cultures from both the nares and the oropharynx were positive for *S. aureus*. Intermittent carrier - between 10% and 80% of cultures were positive from both the nares and the oropharynx. Persistent carrier - $\geq 80\%$ of cultures were positive from either the nares or the oropharynx.

**Table 5:**
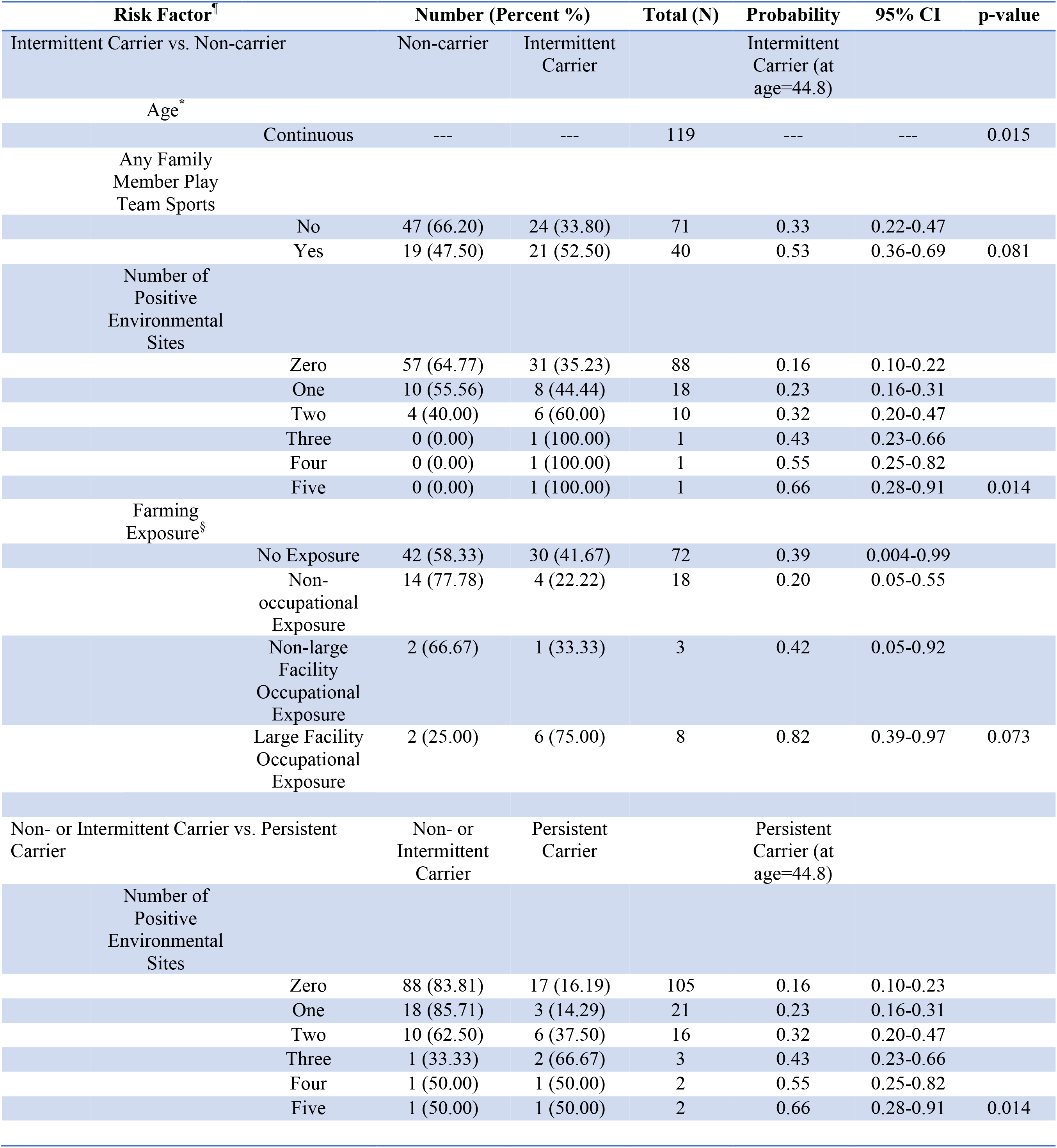
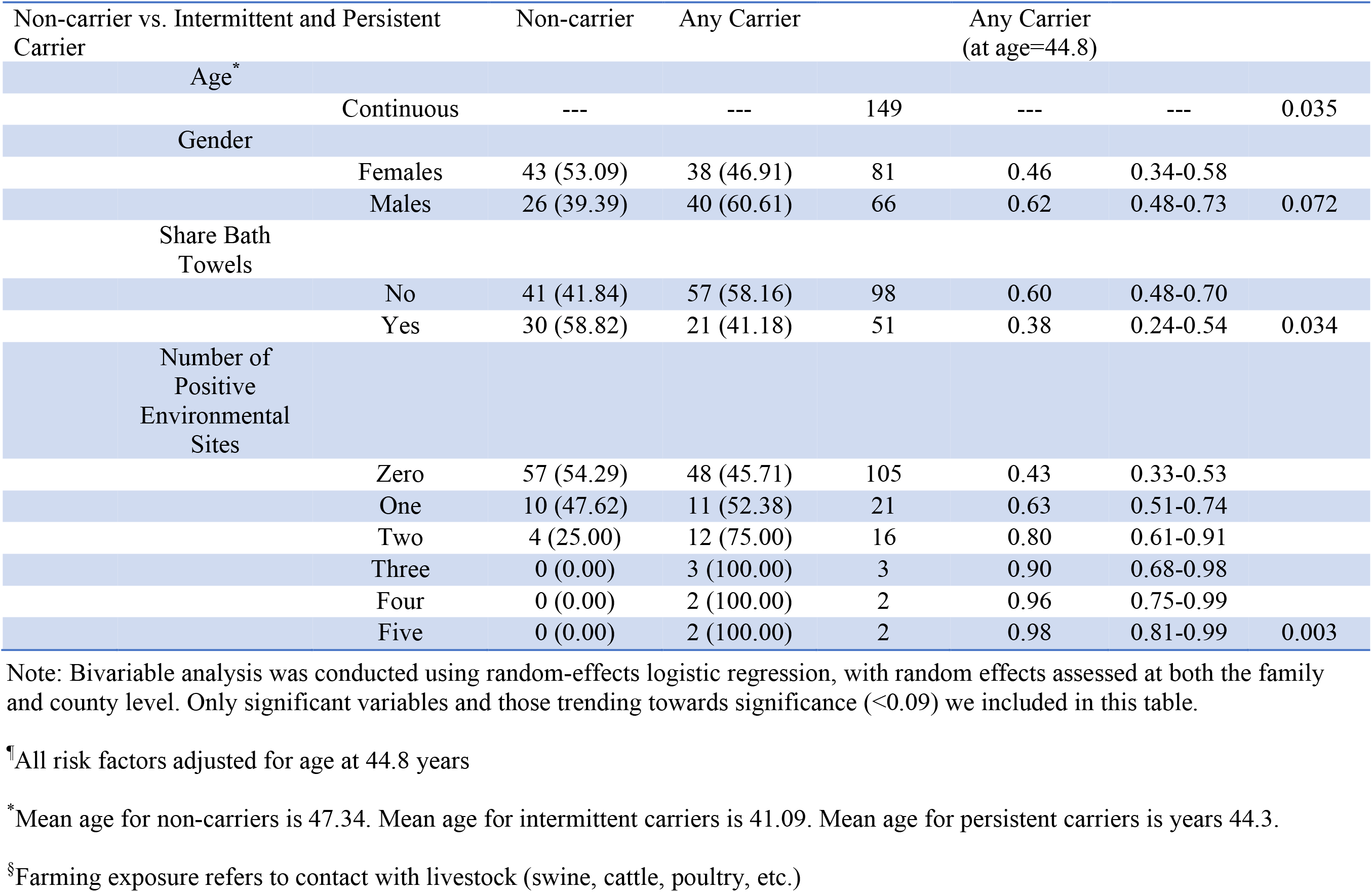
Trending and significant participant demographics and risk factors

**Table 6:** Trending and significant participant demographics and risk factors for minors

| Risk Factor<sup>¶</sup> |  | Number (Percent %) |  | Total (N) | Probability | 95% CI | P-value |
| --- | --- | --- | --- | --- | --- | --- | --- |
| Non-carriers vs. Intermittent Carriers |  | Non-carrier | Intermittent Carrier |  | Intermittent Carrier (at age=9.65) |  |  |
| Race |  |  |  |  |  |  |  |
|  | Caucasian | 29 (50.90) | 28 (49.10) | 57 | --- | --- |  |
|  | Other | 6 (100.00) | 0 (0.00) | 6 | --- | --- | 0.033<sup>#</sup> |
| Non- or Intermittent Carrier vs. Persistent Carrier |  | Non- or Intermittent Carrier | Persistent Carrier |  | Persistent Carrier (at age=9.65) |  |  |
| Age<sup>*</sup> |  |  |  |  |  |  |  |
|  | Continuous | --- | --- | 74 | --- | --- | 0.033 |

Table 6 Continued
| **Number of Positive Environmental Sites** |  |  |  |  |  |  |
| --- | --- | --- | --- | --- | --- | --- |
| Zero | 45 (91.84) | 4 (8.16) | 49 | 0.024 | 0.006-0.53 |  |
| One | 7 (77.78) | 2 (22.22) | 9 | 0.046 | 0.001-0.78 |  |
| Two | 5 (71.43) | 2 (28.57) | 7 | 0.085 | 0.001-0.926 |  |
| Three | 2 (100.00) | 0 (0.00) | 2 | 0.15 | 0.001-0.96 |  |
| Four | 3 (75.00) | 1 (25.00) | 4 | 0.26 | 0.005-0.961 |  |
| Five | 1 (33.33) | 2 (66.67) | 3 | 0.40 | 0.014-0.97 | 0.041 |

Note: Bivariable analysis was conducted using random-effects logistic regression, with random effects assessed at both the family and county level. Only significant variables and those trending towards significance ( $<0.09$ ) we included in this table.
<sup>†</sup>All risk factors adjusted for age (at 9.65 years), gender, and an age-gender interaction term.
\*Mean age for non-carriers is 9.35. Mean age for intermittent carriers is 8.80. Mean age for persistent carriers is 12.79.
#Fisher's Exact Test used to determine p-value. Probabilities and 95% CIs could not be determined for these risk factors due to small sample sizes.

## Discussion

Most investigators assessing *S. aureus* colonization have utilized data on nasal colonization to assign individual persons to one of three categories with regard to carriage: persistent carriers, intermittent carriers, and non-carriers (4, 23, 24). Investigators who utilize only nasal colonization to assess carrier state could misclassify oropharynx-only carriers as non-carriers. In addition, investigators could misclassify individual persons if the time period over which cultures are obtained is too short. Using the data from this manuscript, we estimate that 9 persistent carriers and 23 intermittent carriers (32/154, 20.8%) would have been misclassified if nasal cultures alone were used to identify *S. aureus* carriage (Figure 1). Unrecognized *S. aureus* carriers may transmit *S. aureus* to other persons and, thus, may ‘render infection-control programs futile’ (25). Additionally, Robinson et al. found that knowing the colonization status of a patient from a prior visit was associated with a reduced 30-day mortality from MRSA bacteremia (26), which the investigators felt might reflect the earlier use of glycopeptides in the group with previous MRSA colonization. Thus, their data support the idea that screening some patients to determine their *S. aureus* colonization status may improve outcomes in some patient Populations.

All cultures from 19 of 152 adults (12.5%) assessed in this study were negative for *S. aureus*. However, most (87.5%) participants submitted at least one sample that grew *S. aureus*, indicating most (if not all) people could be colonized. A person’s colonization status may depend upon the extent of contamination in their environment, or on exposure to persons carrying *S. aureus*, rather than the person’s biological susceptibility to the pathogen. The number of positive environmental sites, and farming exposure, (which nearly met the 0.05 significance level) both assess environmental pressures and level of exposure to *S. aureus*. The number of contaminated environmental sites was significantly associated with *S. aureus* carriage in all analyses for adults, and with being an intermittent carrier when compared with non-carriers for minors. Both of these variables support the hypothesis that that given adequate exposure, any person could be colonized.

Van Belkum et al. inoculated *S. aureus* into participant’s nares and observed that those classified as non-carriers and intermittent carriers lost *S. aureus* colonization on average 4 and 14 days post colonization event, respectively (21). Their results suggest that persons we classified as intermittent or non-carriers may have acquired and cleared *S. aureus* between study samplings. We also found that non-carriers had the longest time without colonization, but also had the fewest colonization events, suggesting that environmental pressure for these participants was low.

Given our definitions, non-carriers must have infrequent colonization events and a short duration of colonization for each event, and persistent carriers must have extended durations of colonization, and fewer colonization events. Of interest, however, is the number of colonization events and average duration of colonization for intermittent carriers. Van Belkum et al. previously showed that non-carriers and intermittent carriers have similar levels of antistaphylococcal antibodies, and that the duration of colonization is not statistically different for the two groups following nasal challenge with *S. aureus* (21). However, in our study, the average duration of colonization for intermittent carriers and non-carriers were significantly different for both adults and minors (Table 4). A distribution demonstrated that the duration of colonization for a majority of participants was clustered between 0 and 25 days, which suggests that a majority of intermittent carriers may be similar to non-carriers (Figure S3). In contrast, some intermittent carriers had an average colonization duration longer than 25 days and may be similar to persistent carriers. For example, one participant had 3 colonization events with an average duration of 85.8 days. This participant did not meet the definition of persistent carriage because he/she had at least one positive culture at 79% of samplings, thus not meet the cutoff of ≥80% required to be categorized as a persistent carrier. Thus, our results suggest that the participants we categorized as intermittent carriers may be a mixed population, some whose carriage dynamics are more like non-carriers and some whose colonization dynamics are more like persistent carries.

Only one swab obtained from putative infections grew *S. aureus* when cultured. This sample was the only one collected by a participant’s physician, suggesting that participants may have obtained inadequate samples because we did not teach participants to obtain samples from possible infections. Alternative explanations for these results are that these participants had infections caused by pathogens other than *S. aureus*, or that they had non-infectious skin lesions.

The observed oxacillin resistance levels using genotypic and phenotypic assays are similar for both unadjusted (2.7 vs. 2.07) and adjusted methodologies (5.4 vs. 4.71), indicating the subset of isolates submitted for MIC testing was likely representative of the entire set of 3,185 isolates. Overall, the levels of antimicrobial resistance for the adjusted isolate set were low, with resistance levels of less than 10% for all but two antimicrobials.

The minimum spanning tree revealed that patterns of *spa* types were similar for the two counties (Figure 2). We did not observe differences amongst the lineages, suggesting the same strains circulate within both populations. This observation is similar to the observation by Melles et al. that the *S. aureus* population structures in the United States and in Denmark are similar (27).

**Figure 2:**
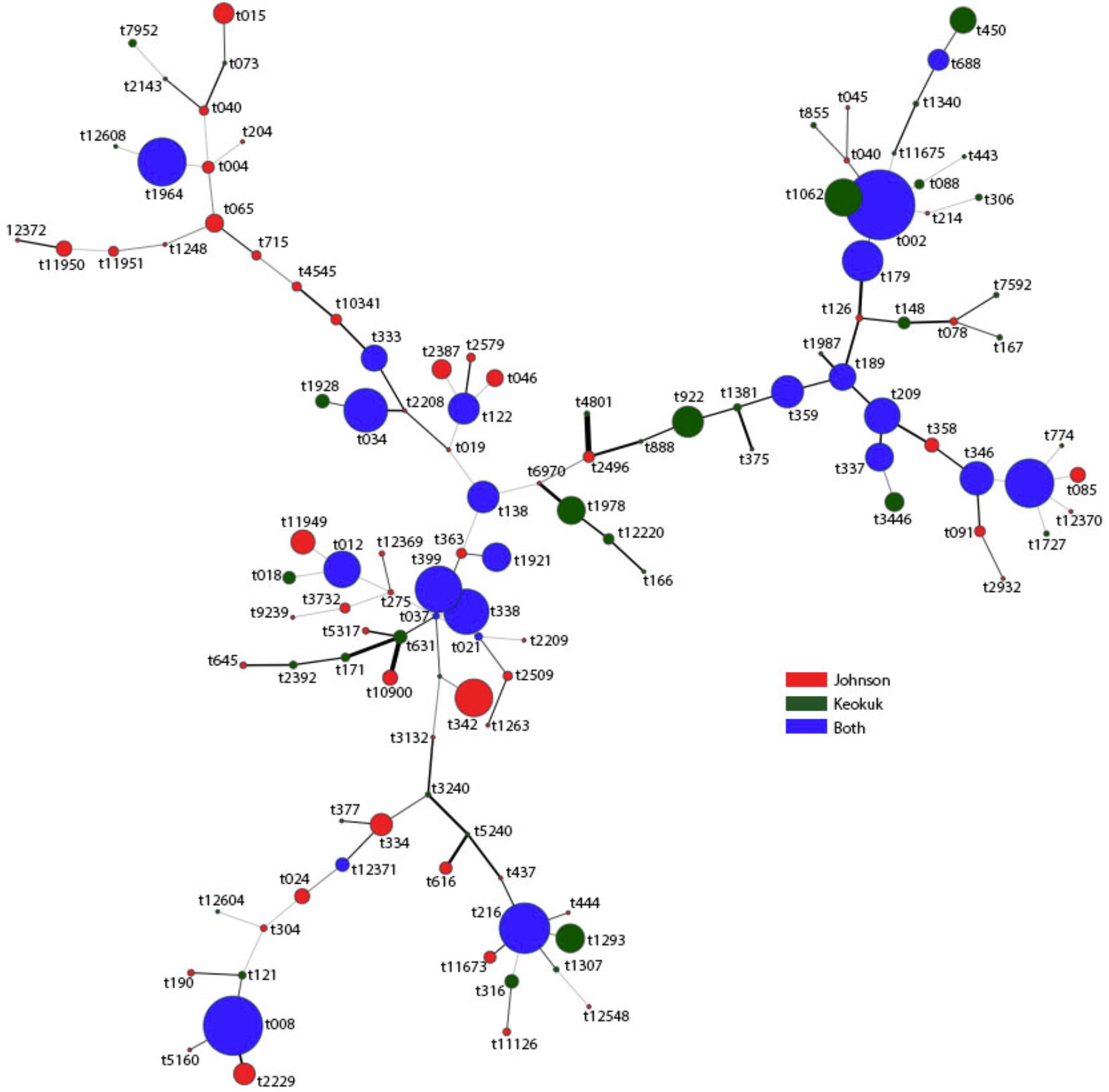
Minimum spanning tree of *spa* type diversity among 147 isolates comprising 57 *spa* types by county. The size of the circle is proportional to the number of isolates within the set; the thickness of connecting lines is proportional to the genetic distance.

**Figure 3:**
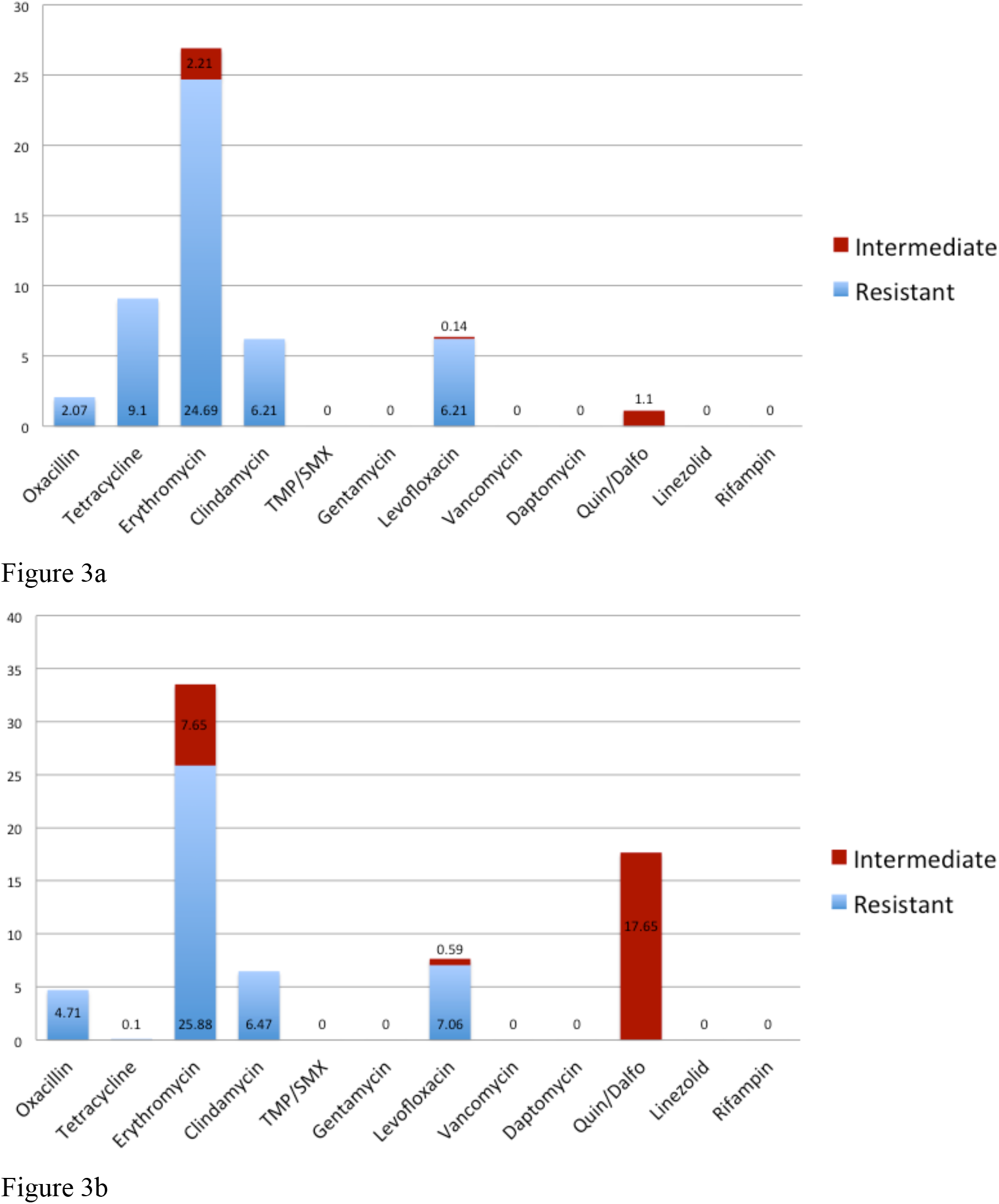
Antimicrobial susceptibility to a panel of antimicrobials tested via minimum inhibitory concentration. A. Unadjusted results (n=725). B. Adjusted results (n=170).

We found a significant inverse relationship of age with intermittent carriage given one is not a persistent carrier. Thus, older participants had a lower probability of being an intermittent carrier. For example, the probability of a 30-year-old participant being colonized (probability: 0.5439) was higher than that a participant who was 60 (probability: 0.2657). We also found a similar inverse association of age and colonization (defined as being an intermittent carrier or persistent carrier).

This study had several strengths. We enrolled family units and we enrolled persons from urban and rural settings, which allowed us to adjust for random variability, and to observe familial transmission events. Longitudinal follow-up, with an average of 34.7 swab sets for adults and 30.9 nasal swabs for minors, allowed us to estimate the frequency of transmission events and of symptomatic infection. Delacour et al. previously showed that *S. aureus* can be viable 18 days after collection in non-charcoal liquid transport medium (28). In addition, we trained participants to swab their nares and oropharynges. Thus, we maximized our ability to detect *S. aureus* colonization and minimized participant inconvenience.

This study had potential limitations. Patients reported symptomatic SSTIs and they obtained wound cultures. Additionally, only one of five patients who submitted wound cultures had a confirmed *S. aureus* infection. Thus, we may have misclassified noninfectious skin lesions as infections or we may have misclassified infections caused by other organisms as *S aureus* infections. However, *S. aureus* is the most common cause of SSTIs seen in emergency departments (29), suggesting our self-reported SSTIs may be classified correctly as *S. aureus* infections.

An additional limitation is that our study duration of up to 52 weeks may not have been adequate to observe colonization and transmission events for persistent carriers. Our follow-up period far exceeds the timeframe observed by van Belkum et al. that colonization duration was 4 days for non-carriers and 14 days for intermittent carrier (21), allowing us to observe multiple colonization and decolonization events for both non-carriers and intermittent carriers. We censored and truncated data when we calculated the duration of colonization so our estimates of duration of colonization would be as accurate as possible while accounting for time between samples. Because we utilized left and right truncation, we most likely underestimated the true duration of colonization but this method minimized the assumptions required for the calculations and allowed us to establish the most accurate estimates of duration of colonization with the data available.

In conclusion, 20.8% of participants would have been misclassified if oropharyngeal samples were excluded, indicating that the oropharynx is an important screening site for persons from the healthy community who present to a healthcare setting and may be at risk of a *S. aureus* infection. We also found that the average duration of colonization were significantly different for intermittent carriers and non-carriers, which is in contrast to the findings by van Belkum et al. (21), However, intermittent carriers may be a mixed population with some people having carriage dynamics more like noncarriers and some having carriage dynamics more like persistent carriers. Further studies are needed to assess the frequency of colonization at other anatomical sites among healthy persons in the community. In addition, a study that uses biologic markers and does not involve decolonization and subsequent challenge with *S. aureus* could help elucidate the differences between non-carriers, intermittent carriers, and persistent carriers.

## Acknowledgements

The authors would like to thank Elizabeth Chrischilles and Kelley Donham for their guidance in the analysis and preparation of this manuscript. This study was funded by a grant from the United States Department of Agriculture (USDA), contract number 2011-67005-30337.

## SUPPLEMENTAL

**Table S1:** Site Specific Prevalence of Nares and Oropharynx Anatomical Sites by County for Adults

|  | **Johnson Co.** | **Keokuk Co.** | **Total** |
| --- | --- | --- | --- |
| **Nares** | 458/2225 (20.6%) | 756/2085 (36.3%) | 1214/4310<br>(28.2%) |
| **Oropharynx** | 694/2226 (31.2%) | 498/2087 (23.9%) | 1192/4313<br>(27.6%) |

**Figure S1:**
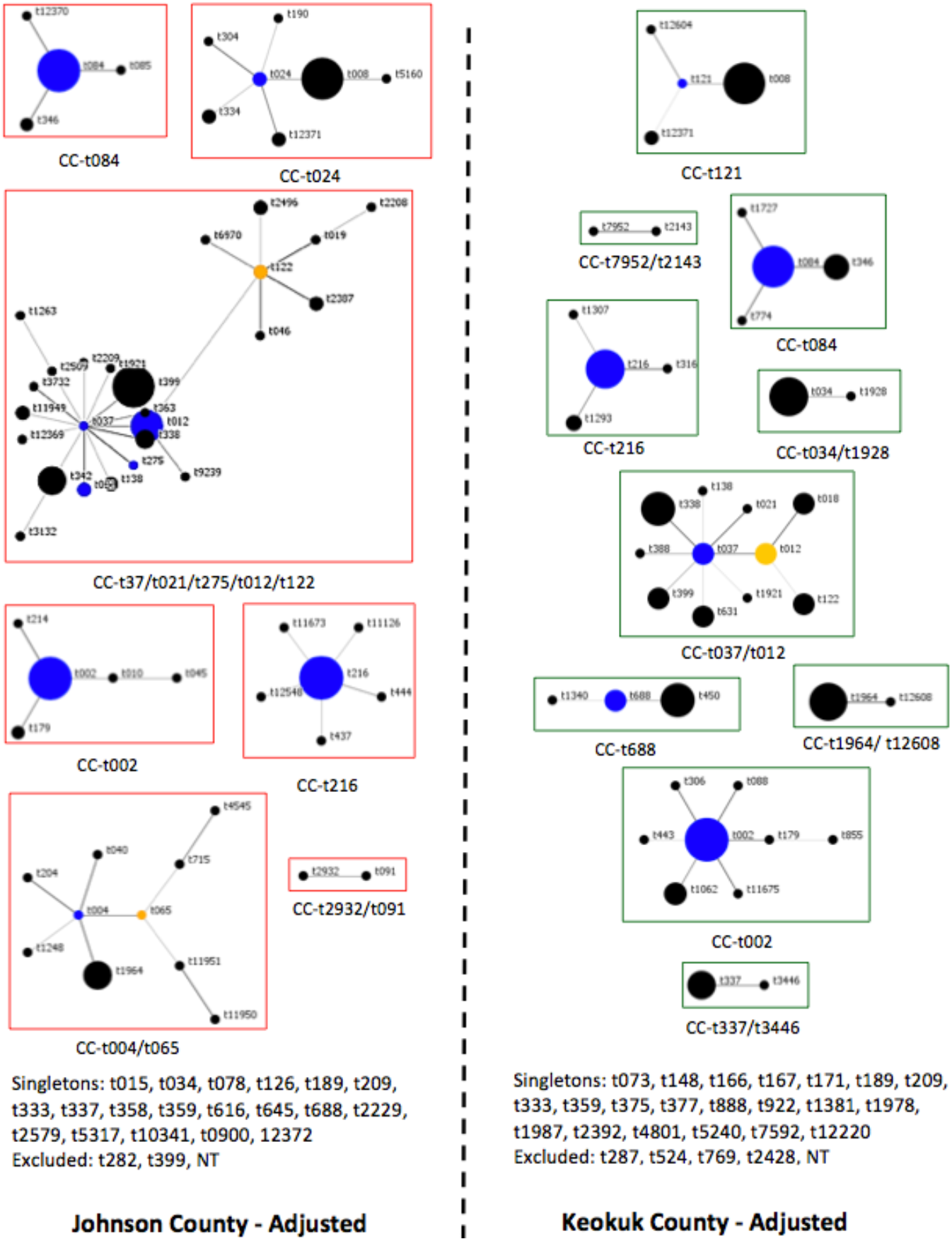
BURP analysis of spa typing data by county, adjusted to control for familial clustering of clonal isolates. Each circle represents a single *spa* type and size of the circle is proportionate to the relative number of isolates of that *spa* type. Blue circles indicate the putative founder, yellow circles are secondary putative founders.

**Figure C2:**
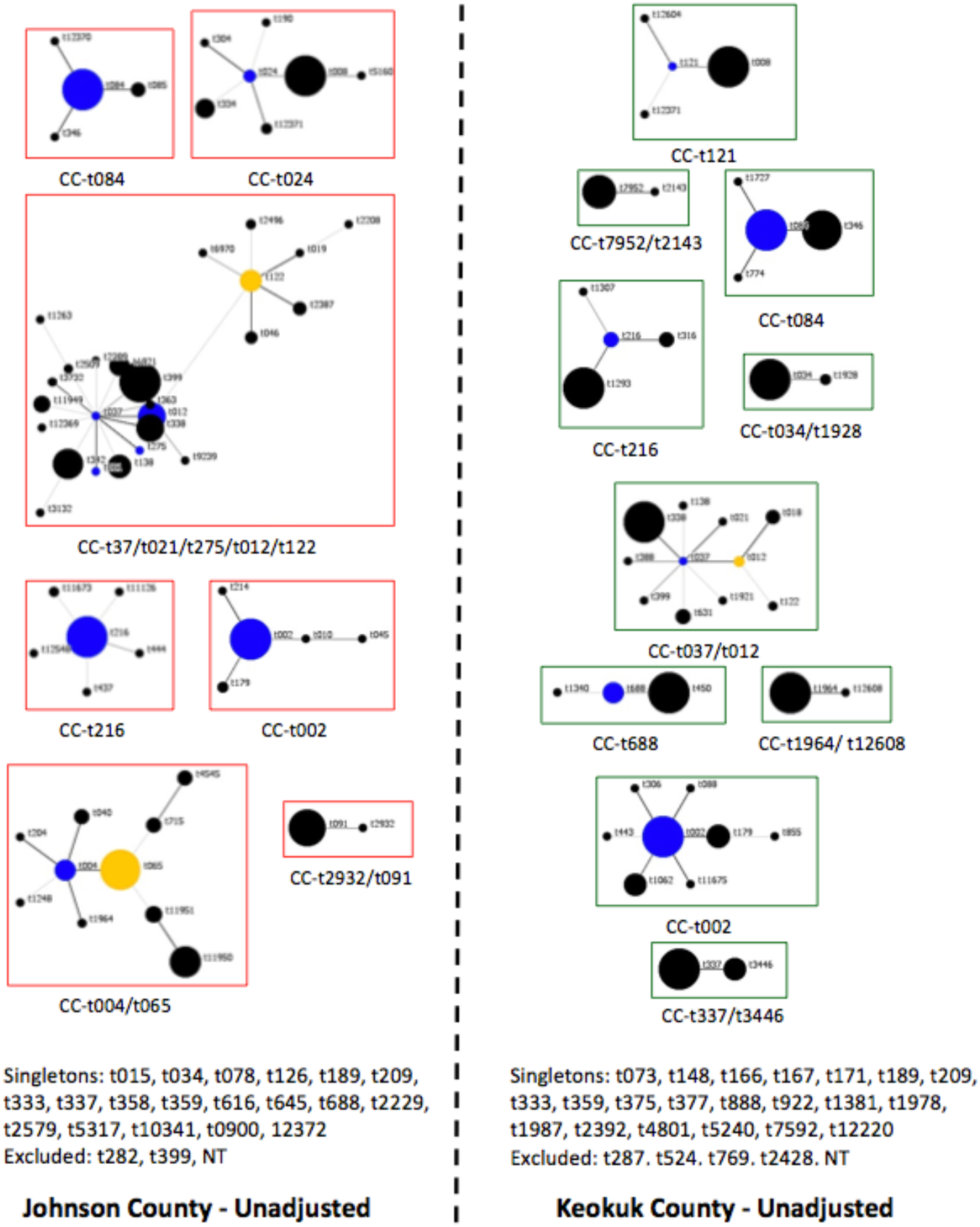
BURP analysis of spa typing data by county. Unadjusted proportions represent all 3184 isolates. Each circle represents a single *spa* type and size of the circle is proportionate to the relative number of isolates of that *spa* type. Blue circles indicate the putative founder, yellow circles are secondary putative founders.

**Figure C3:**
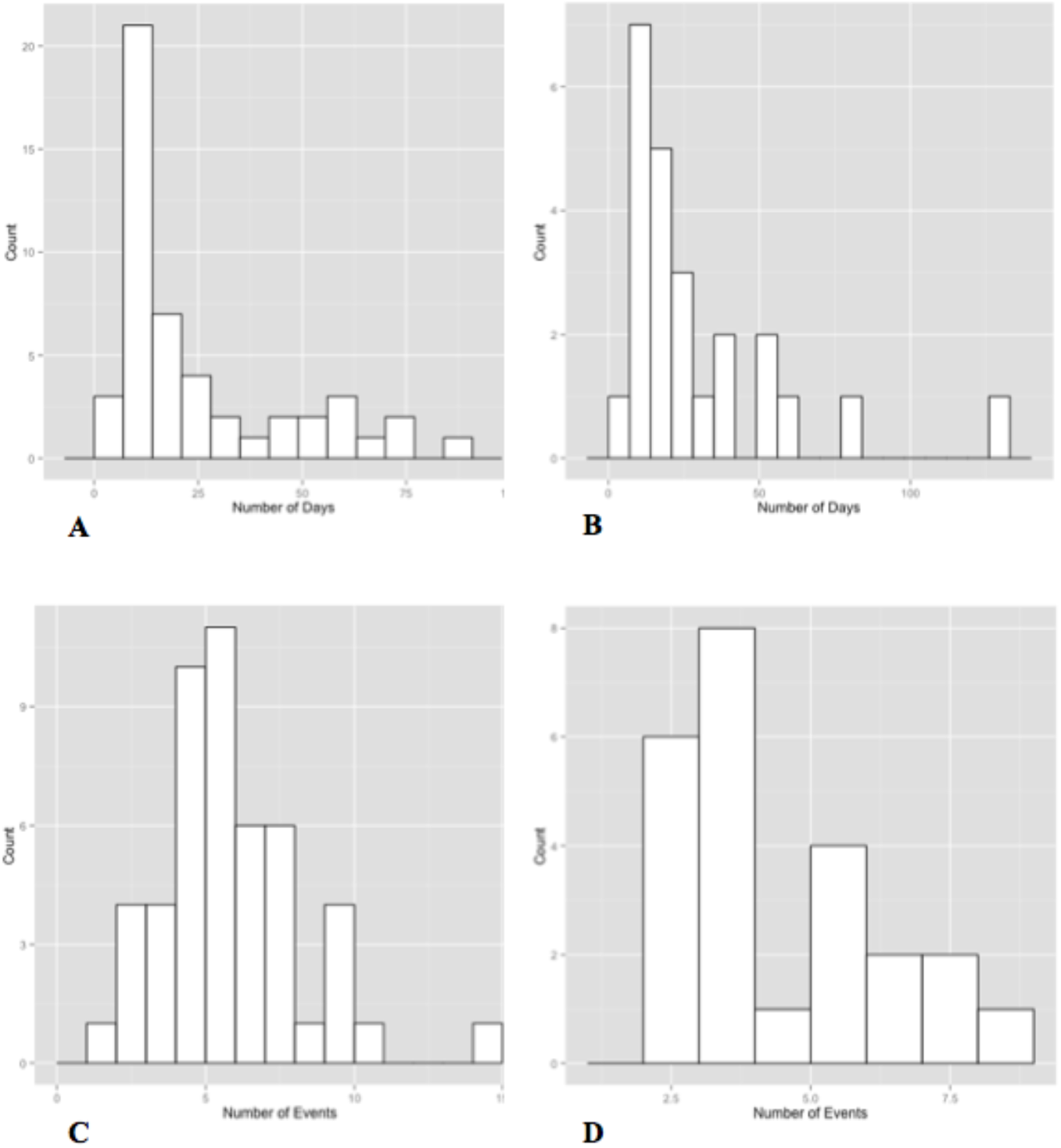
Frequency distributions of the average number of days of colonization and the number of colonization events for intermediate colonizers. A. The average number of days, grouped into 7-day intervals, of adult intermediate colonizers. B. The average number of days, grouped into 7-day intervals, of minor intermediate colonizers. C. The number of colonization events for adult intermediate colonizers. D. The number of colonization events for minor intermediate colonizers.

**Table C2:**
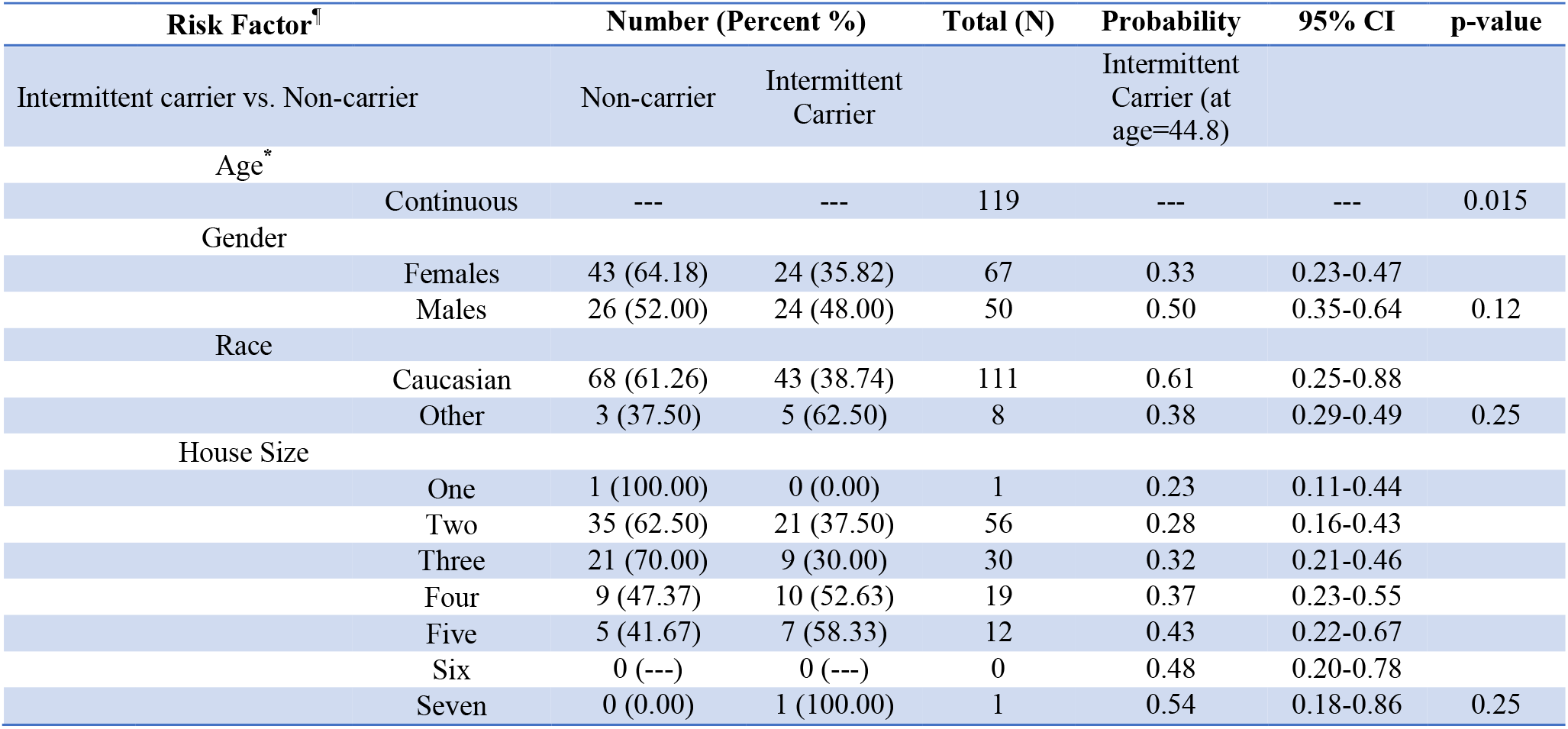
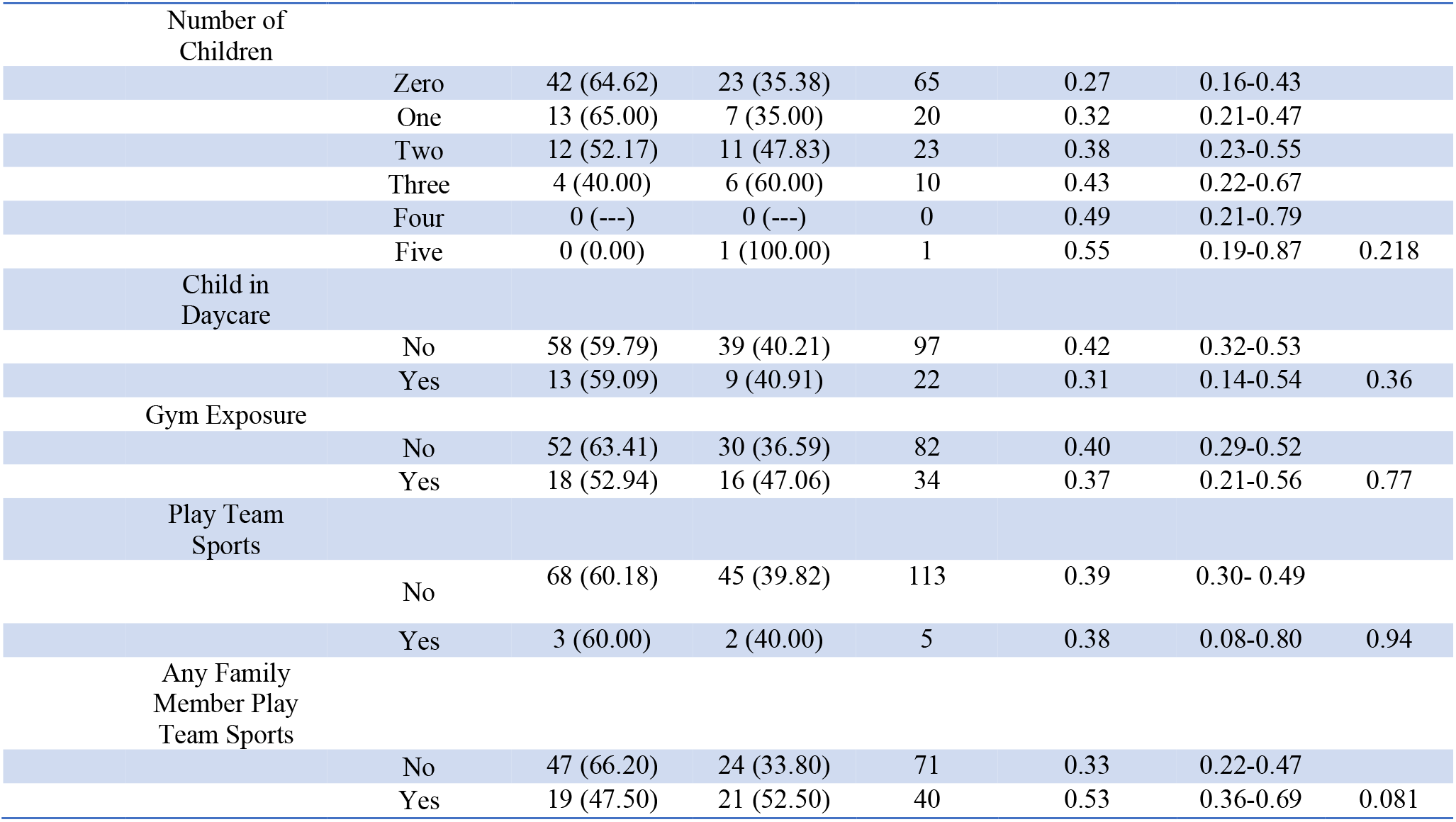
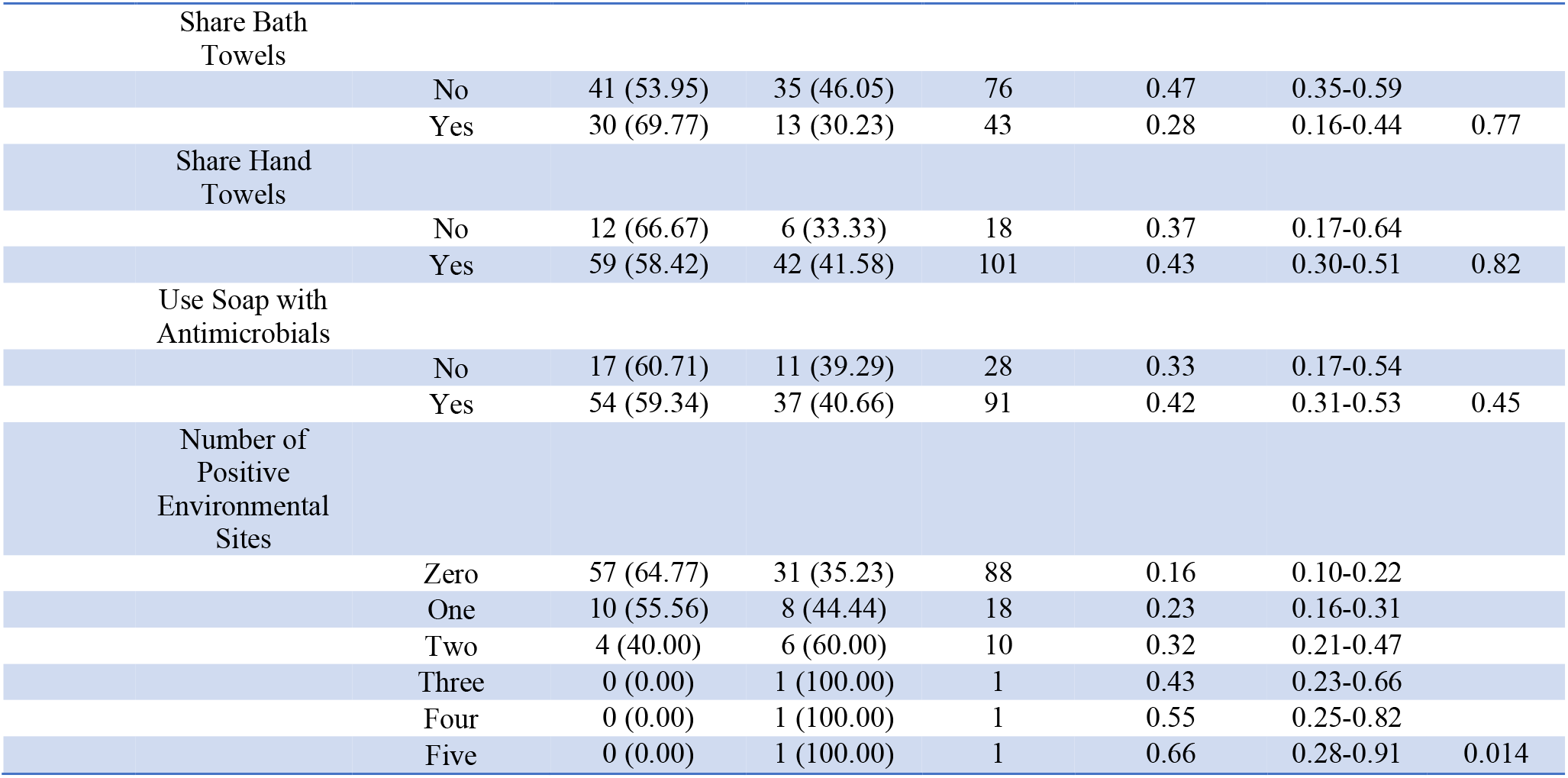
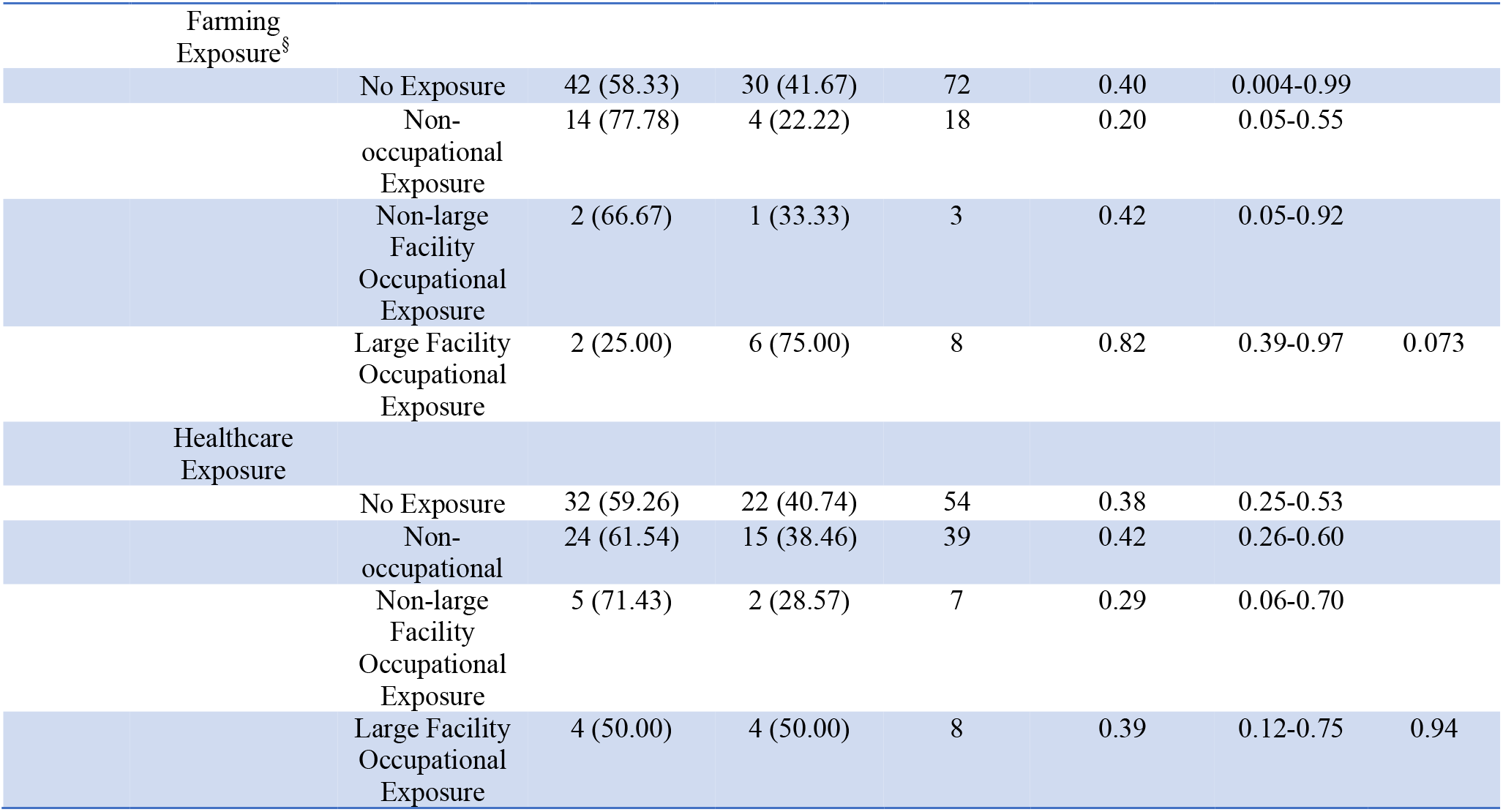
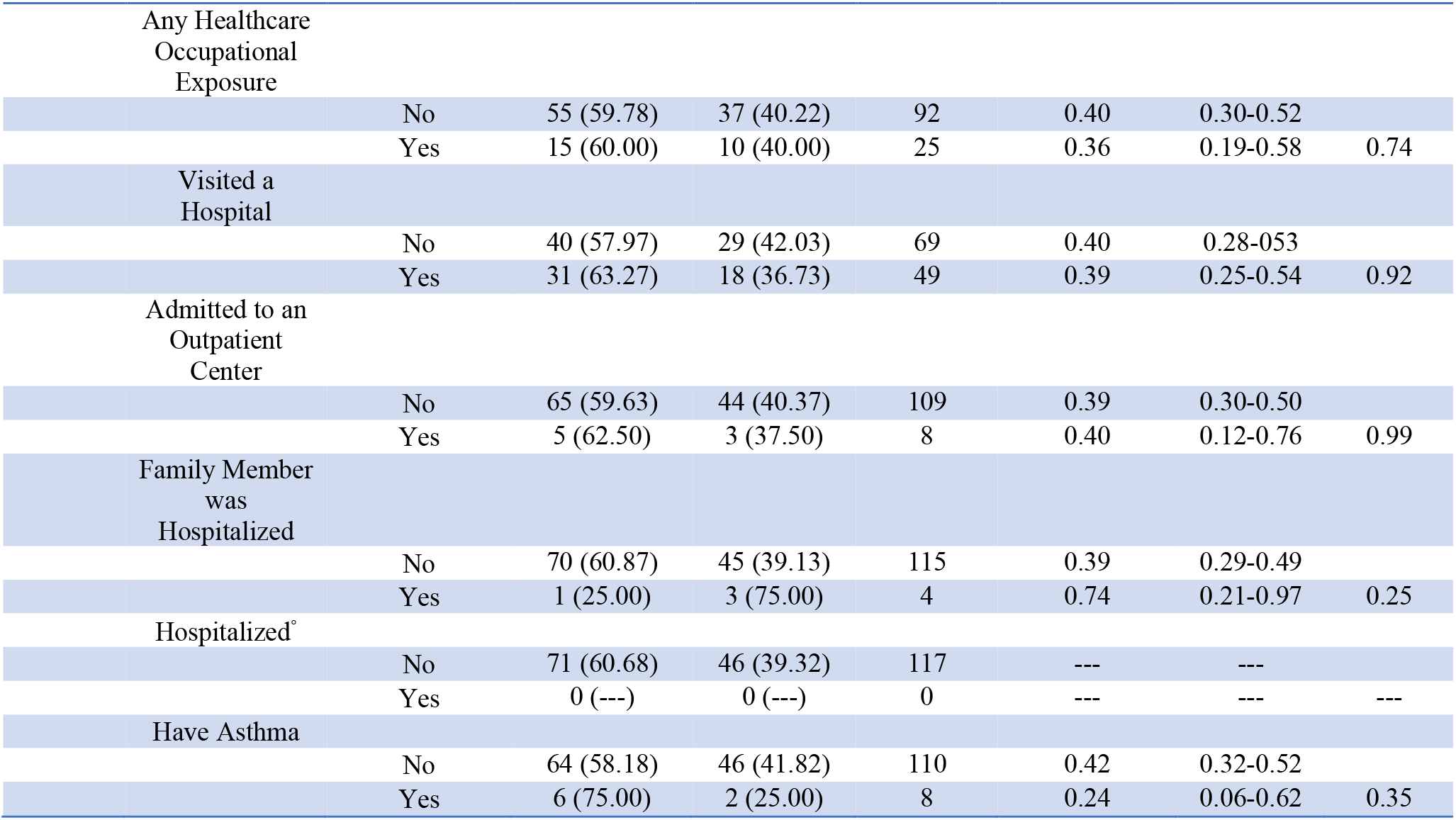
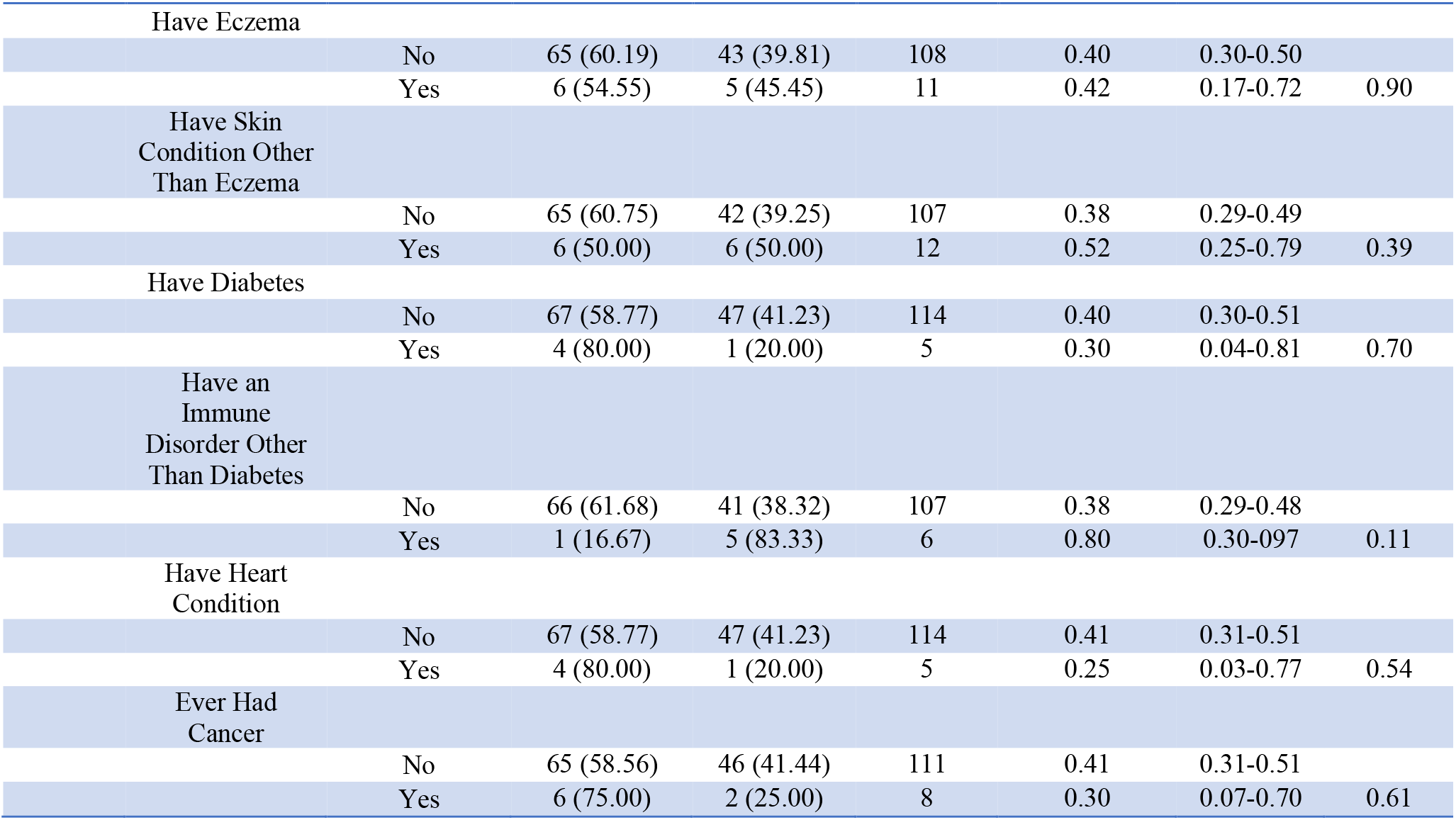
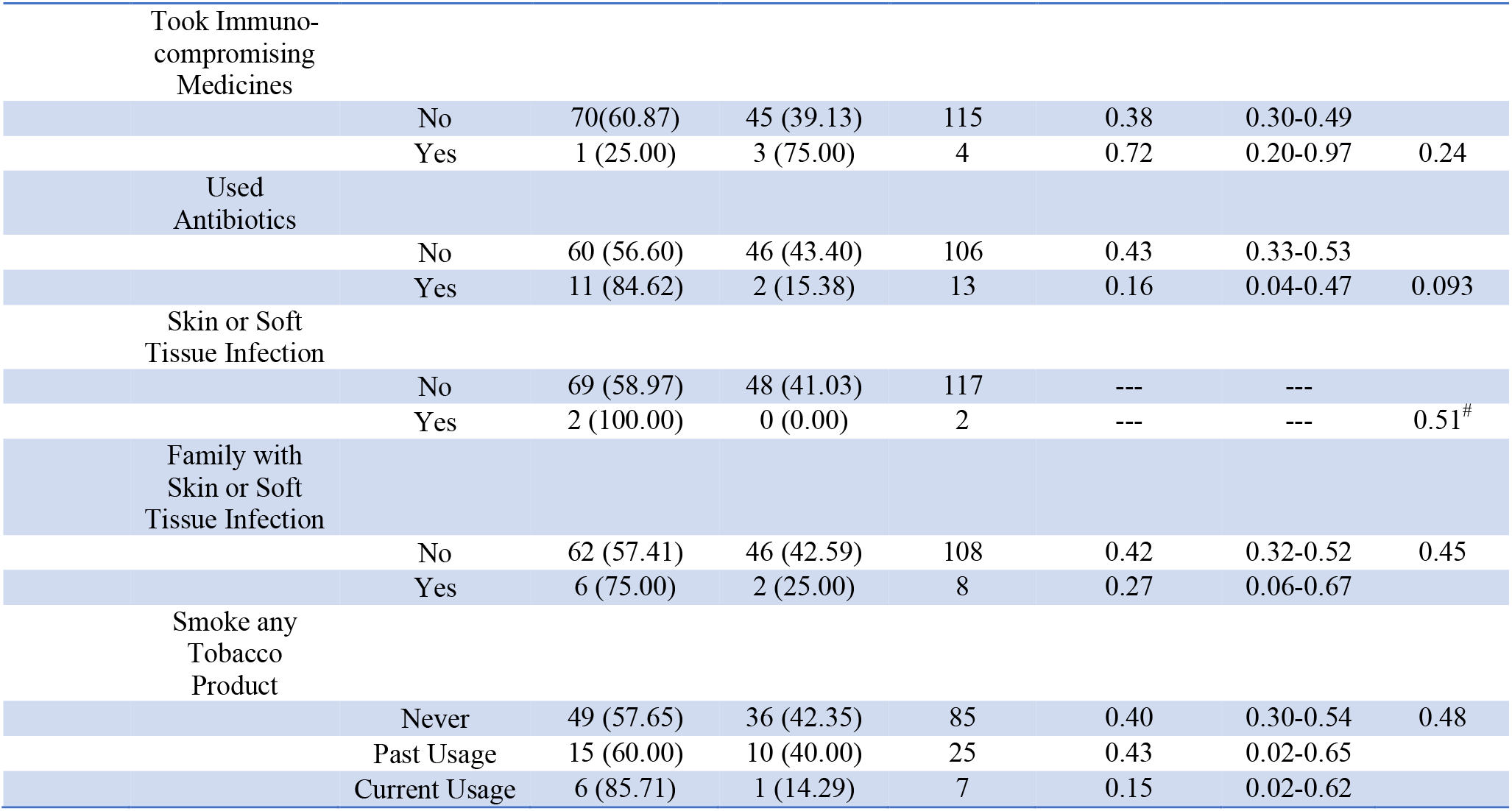
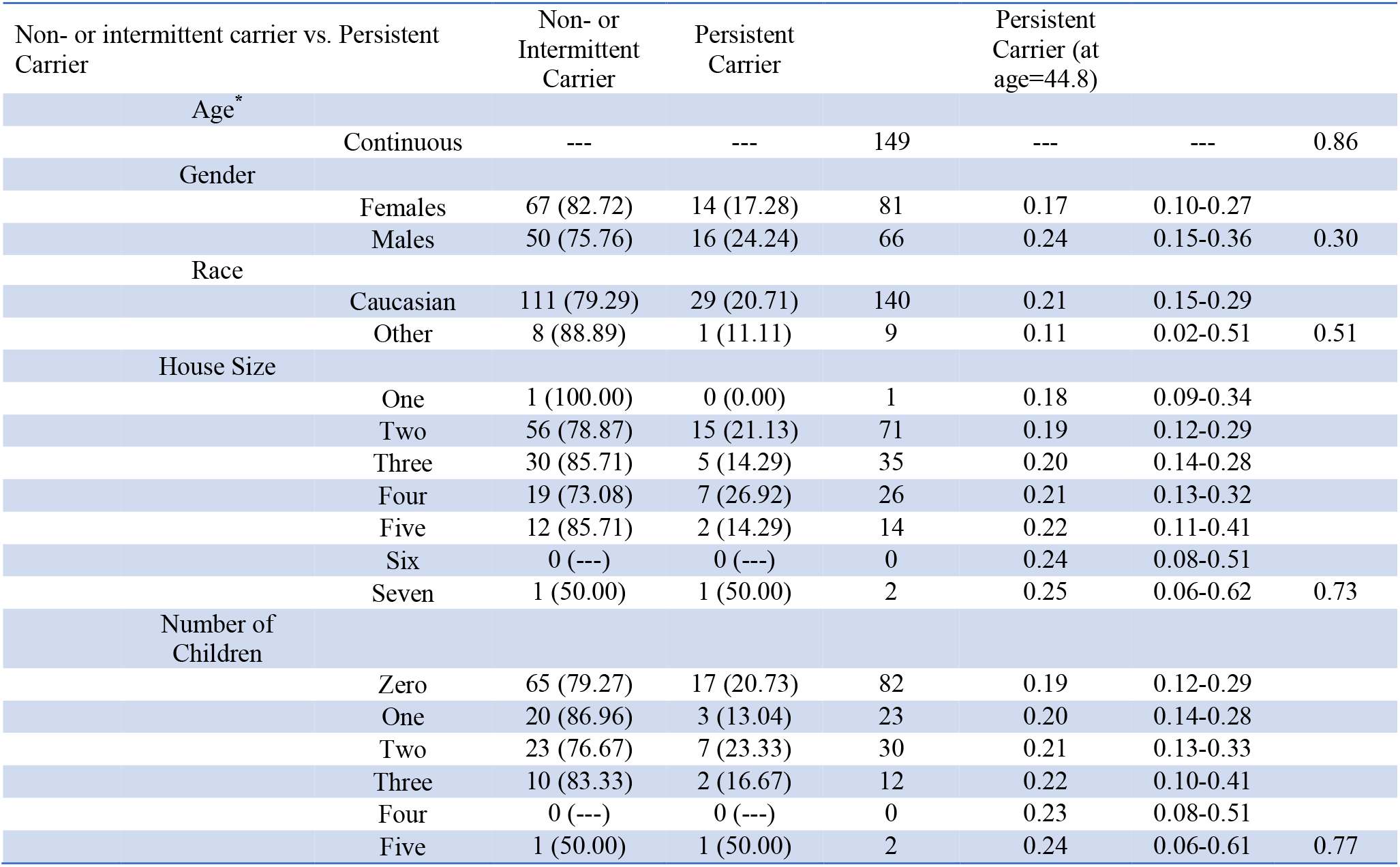
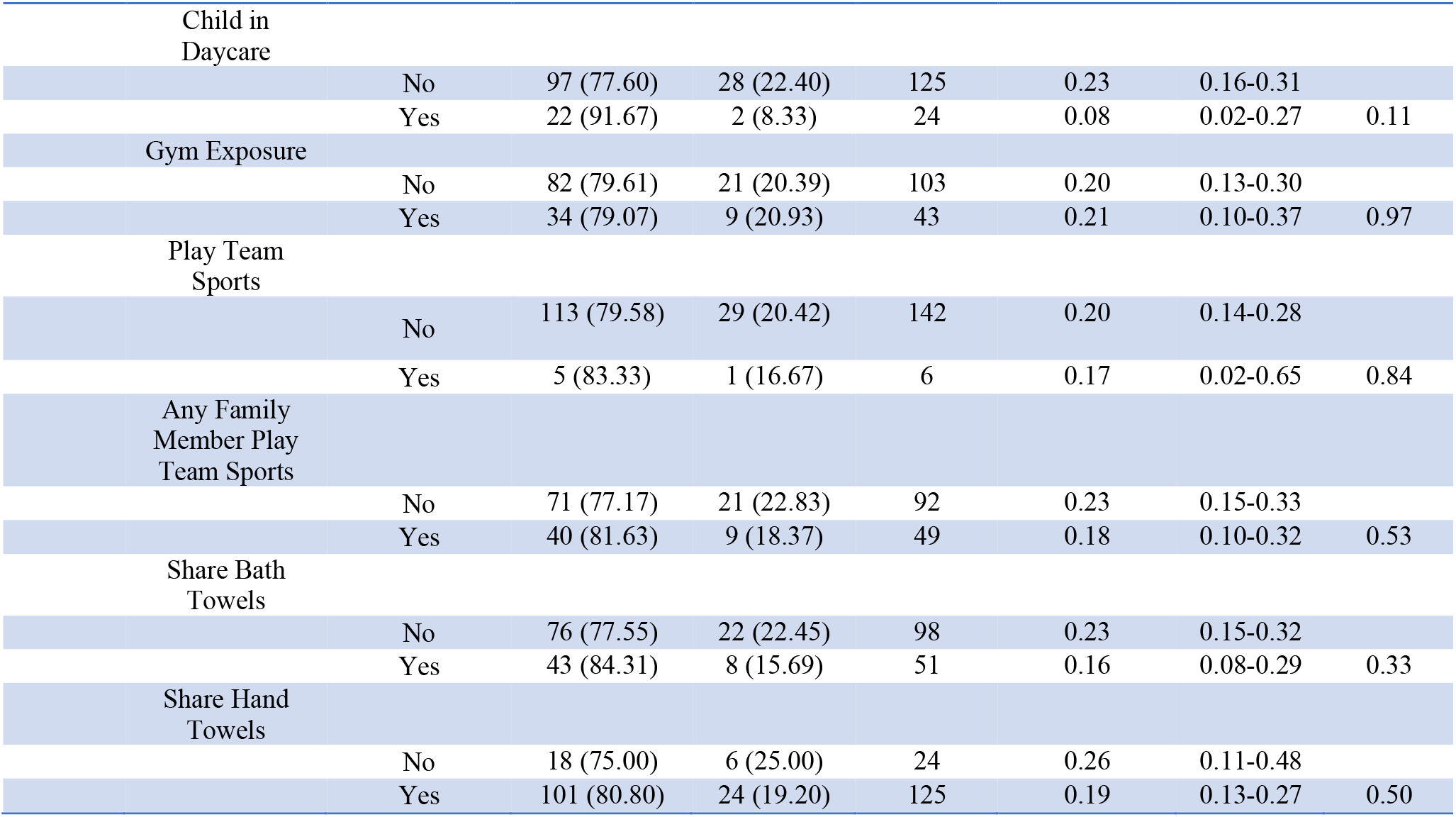
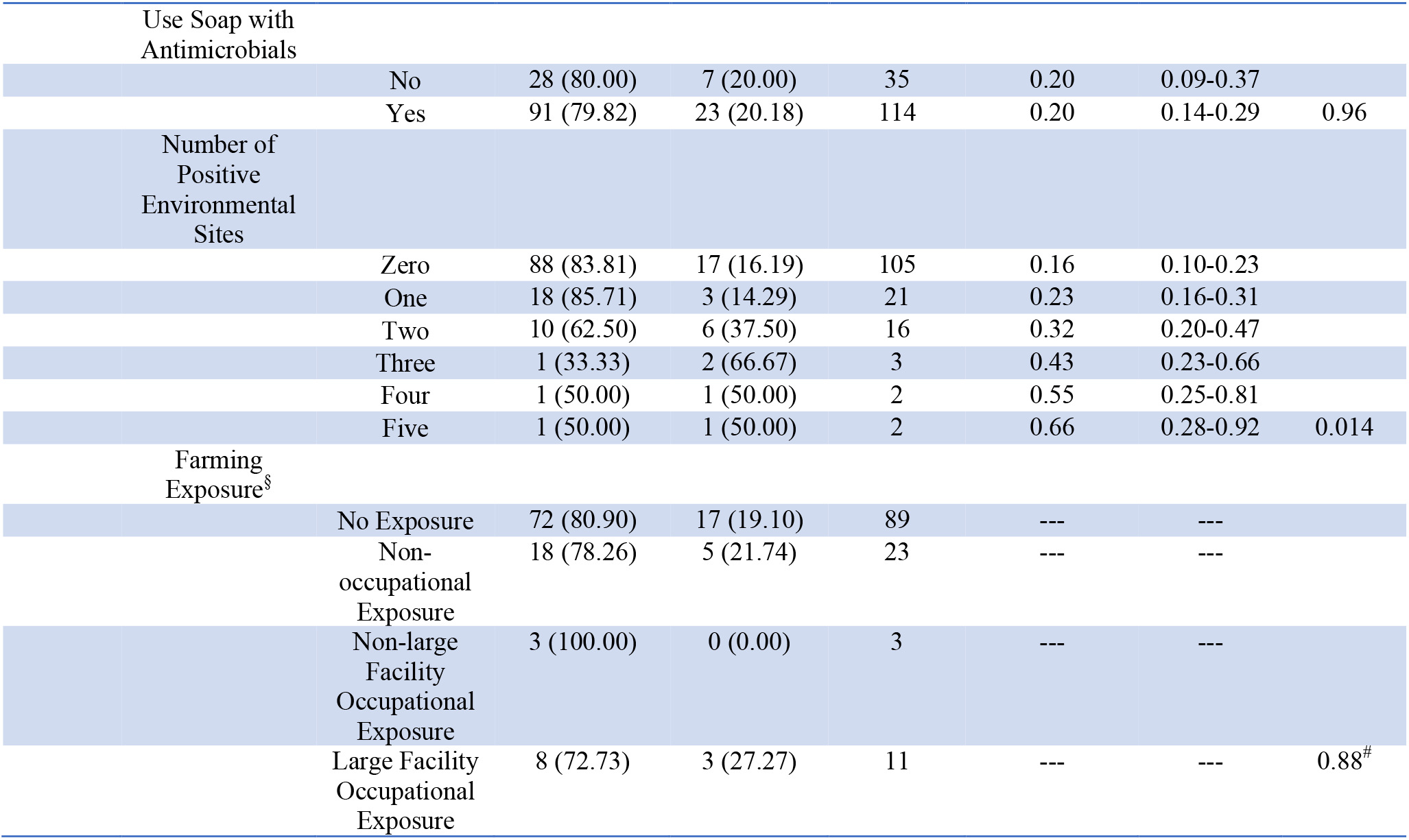
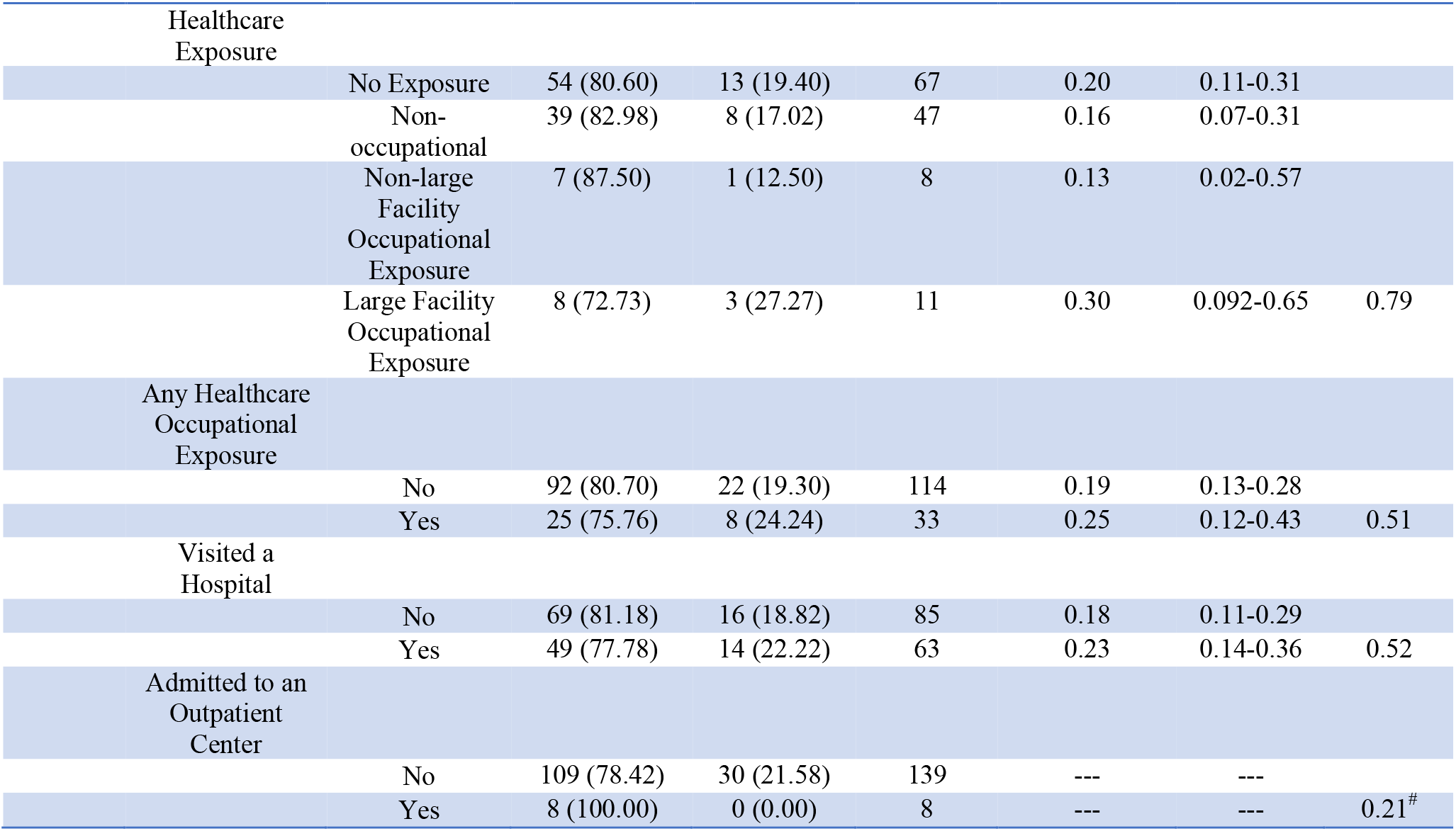
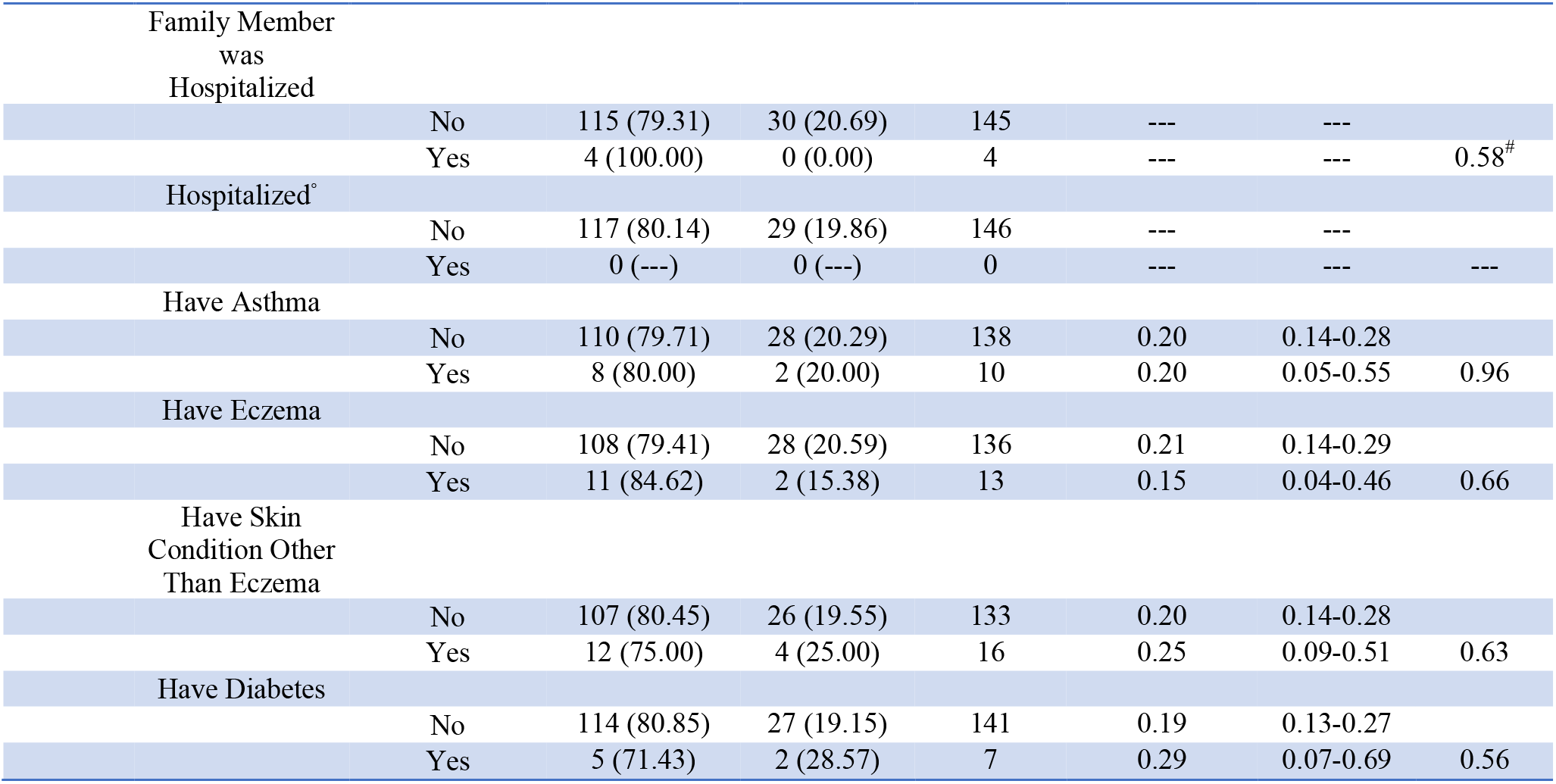
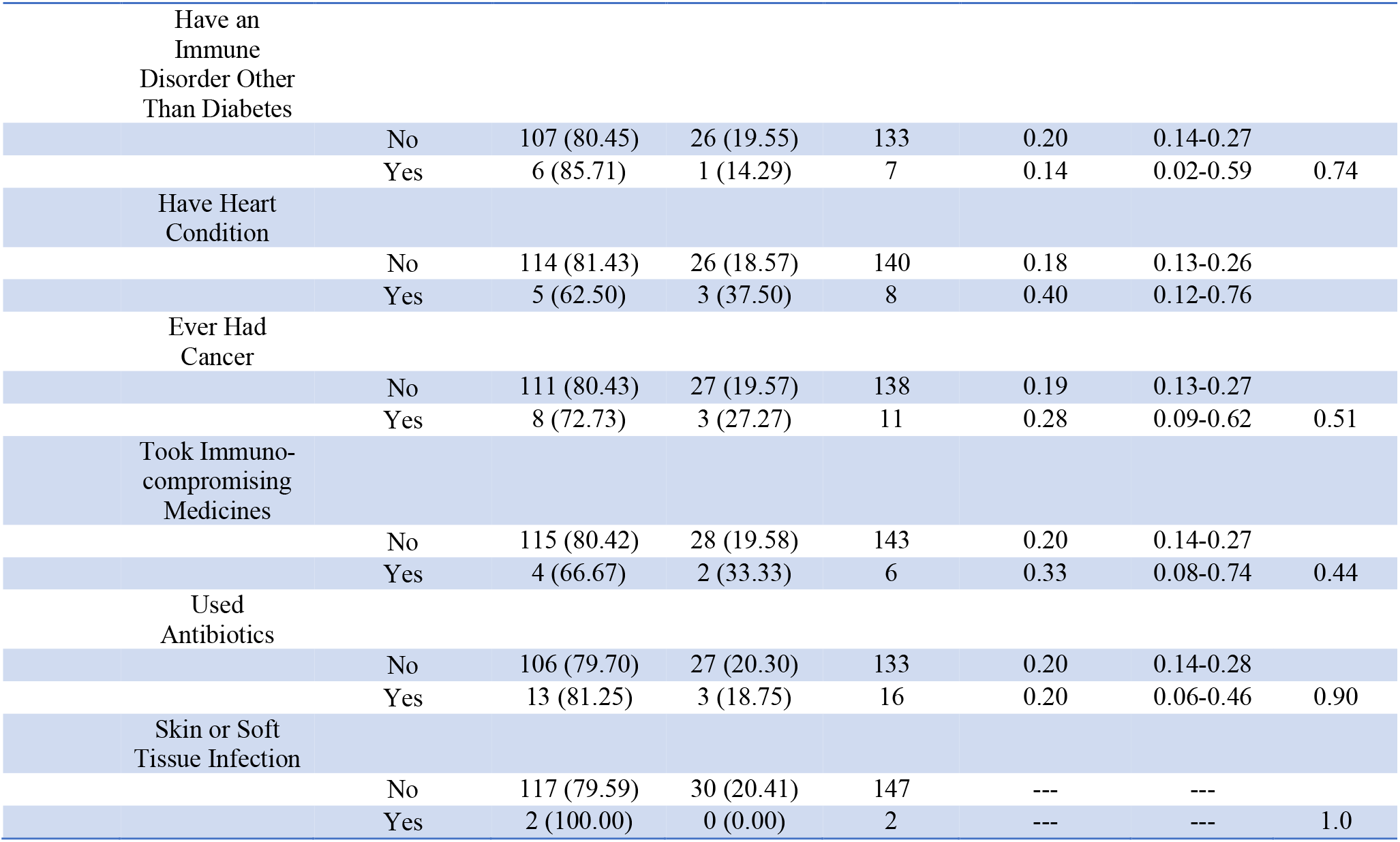
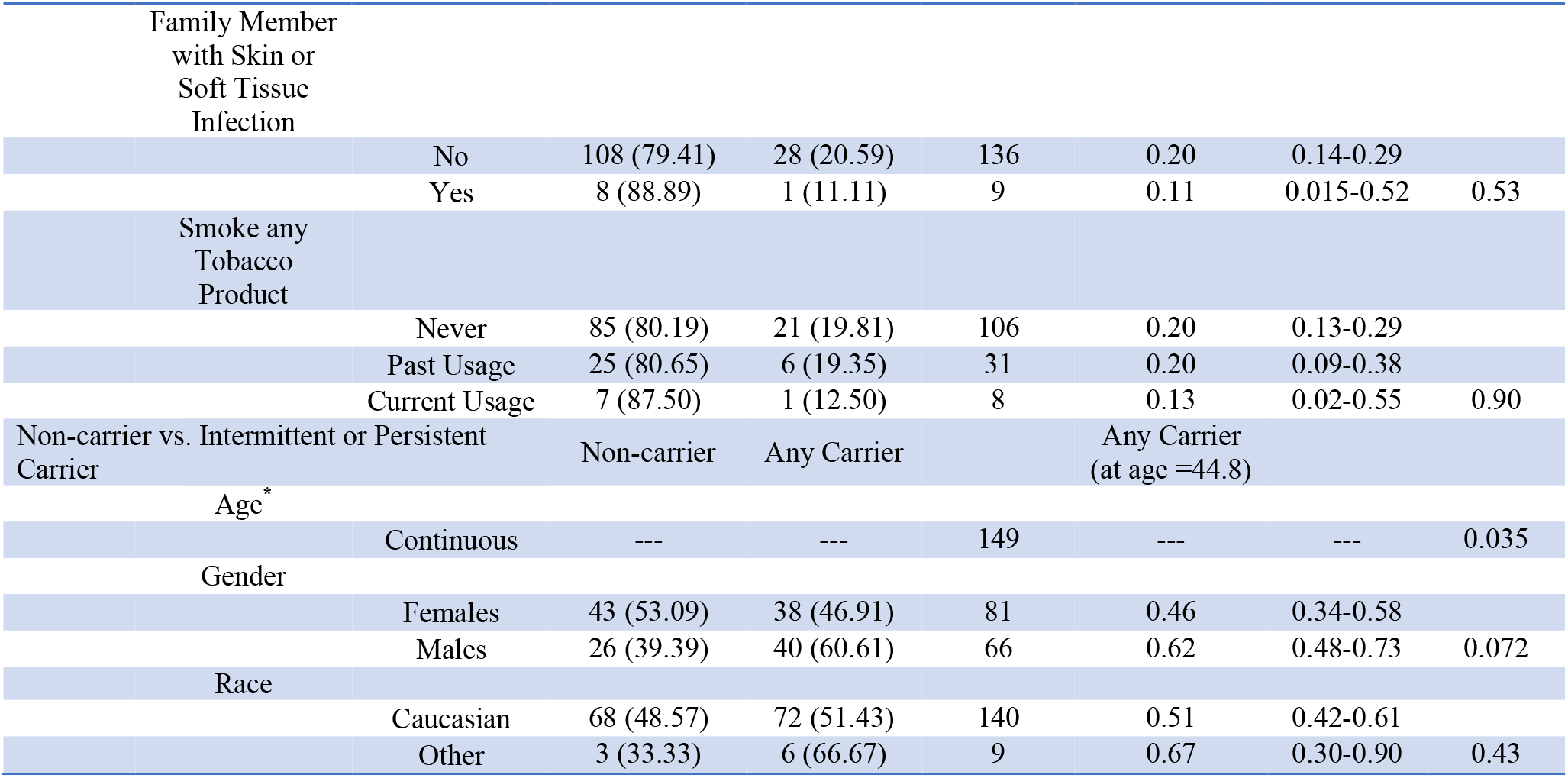
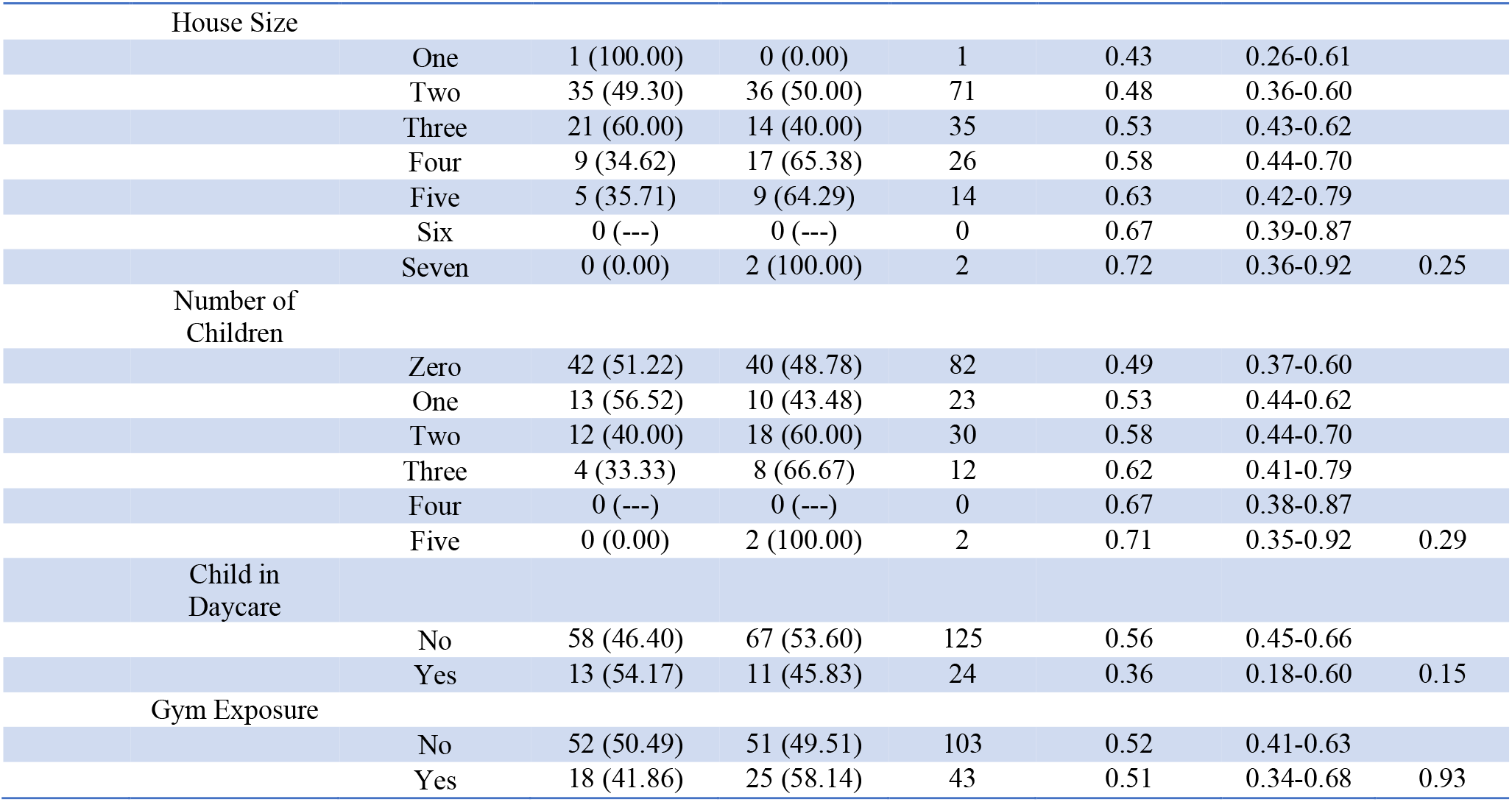
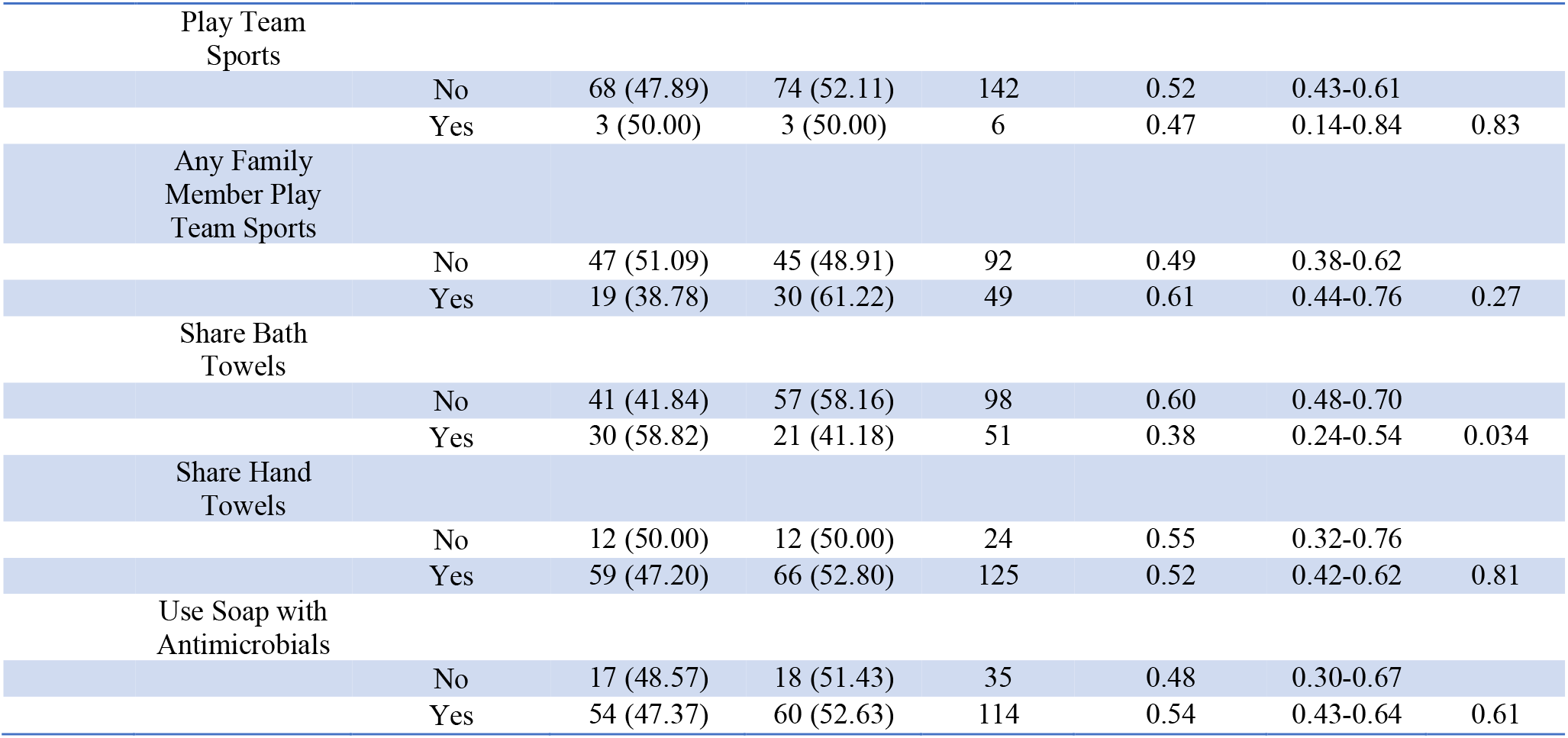
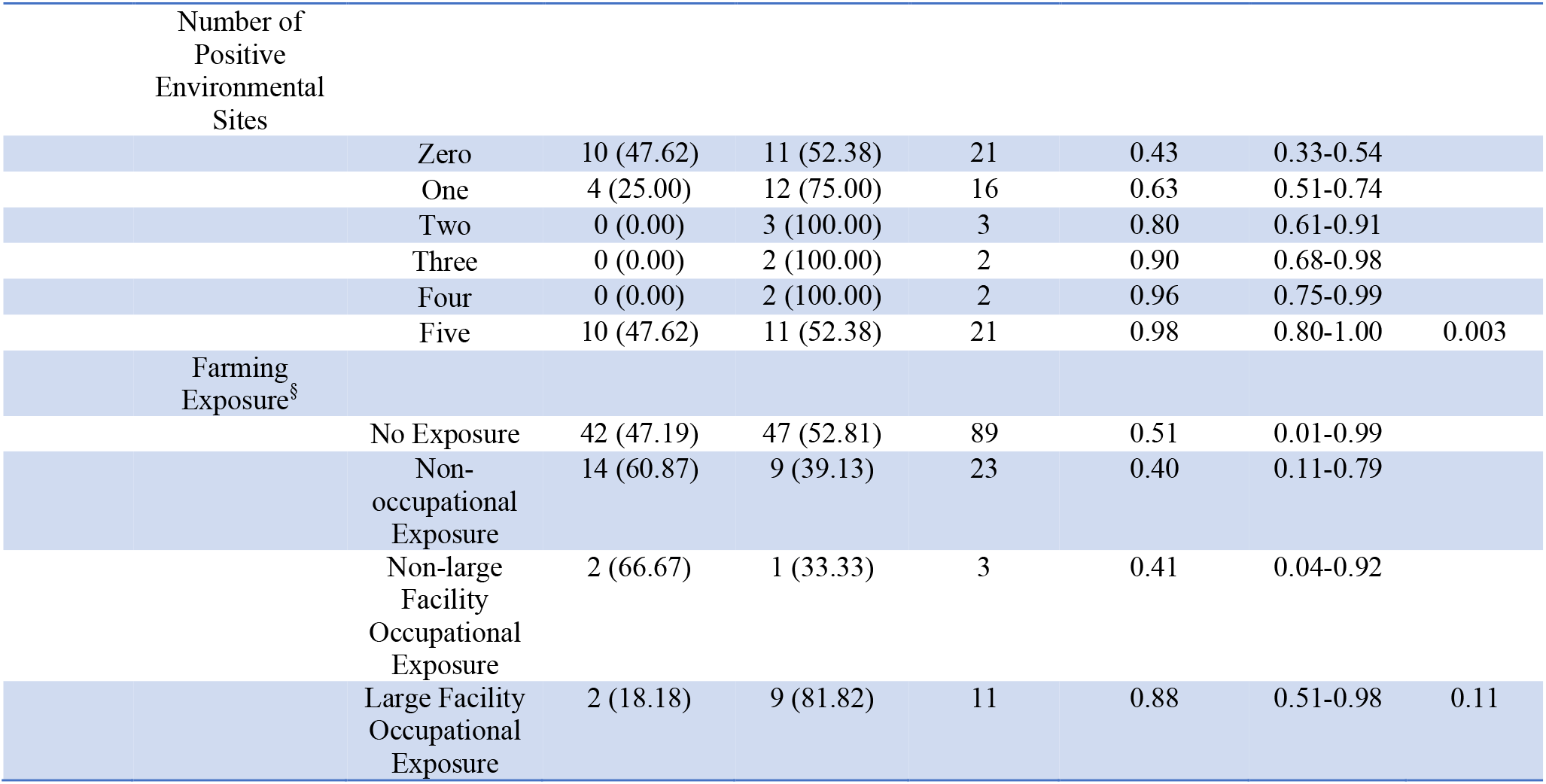
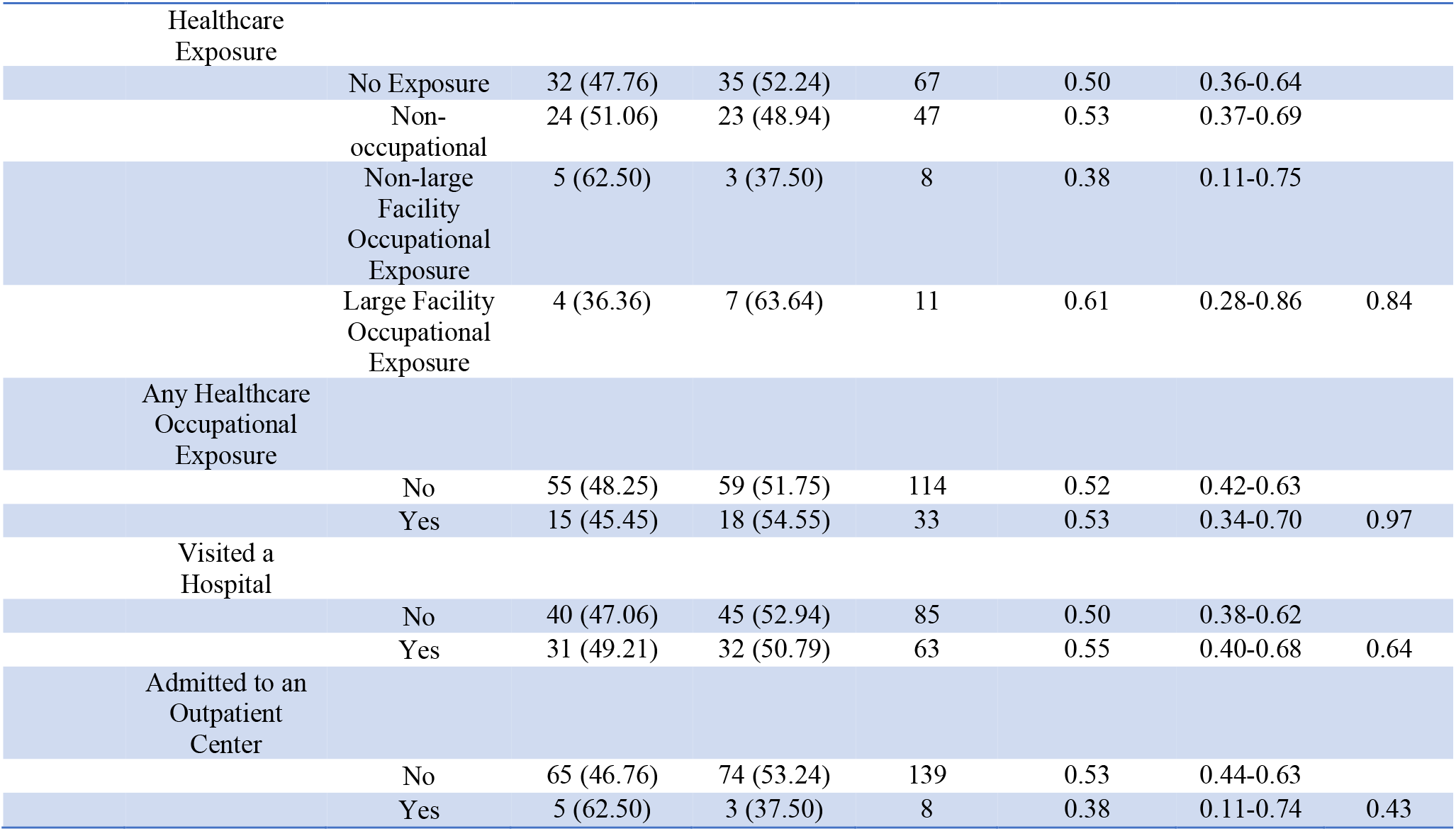
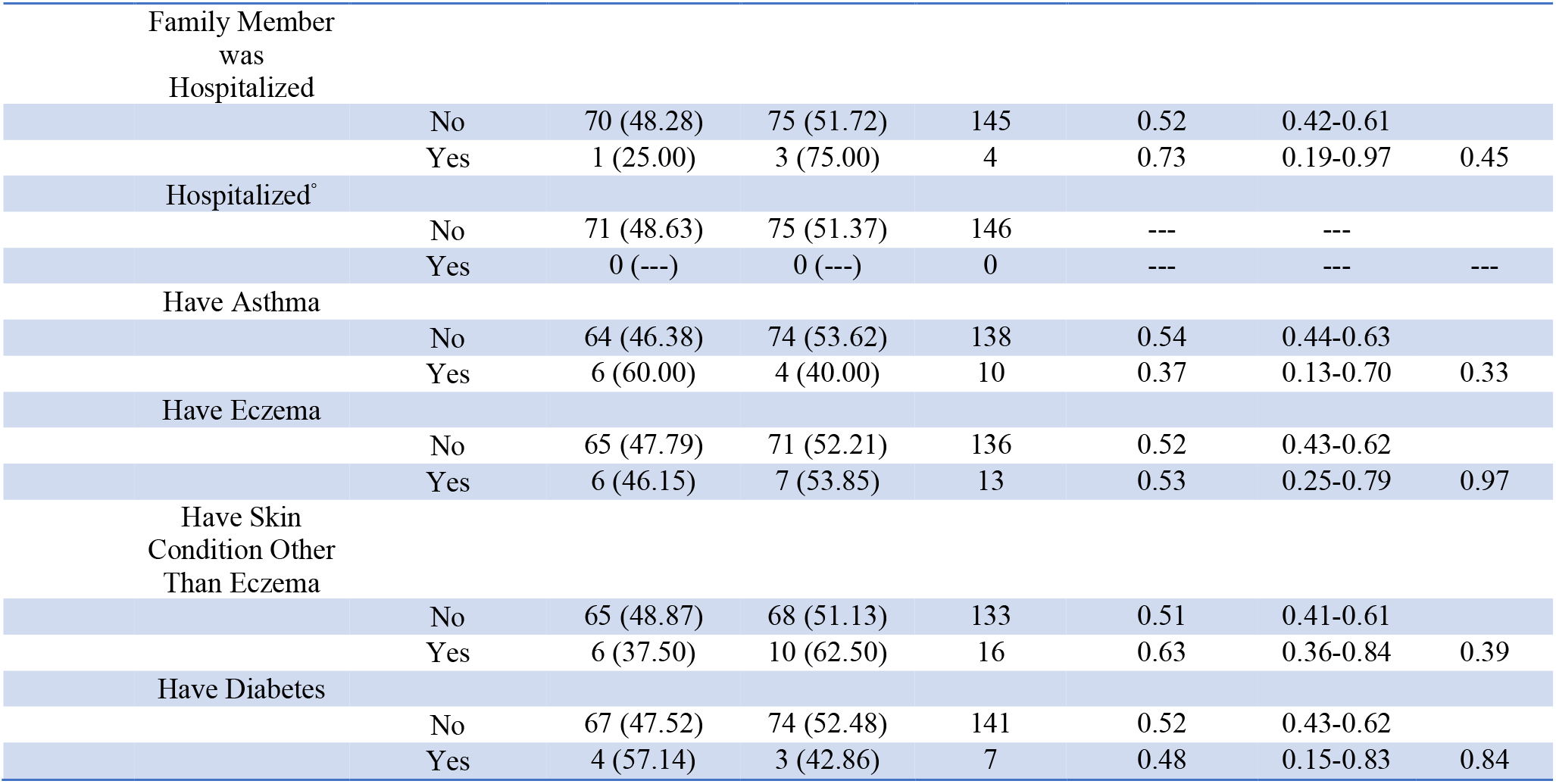
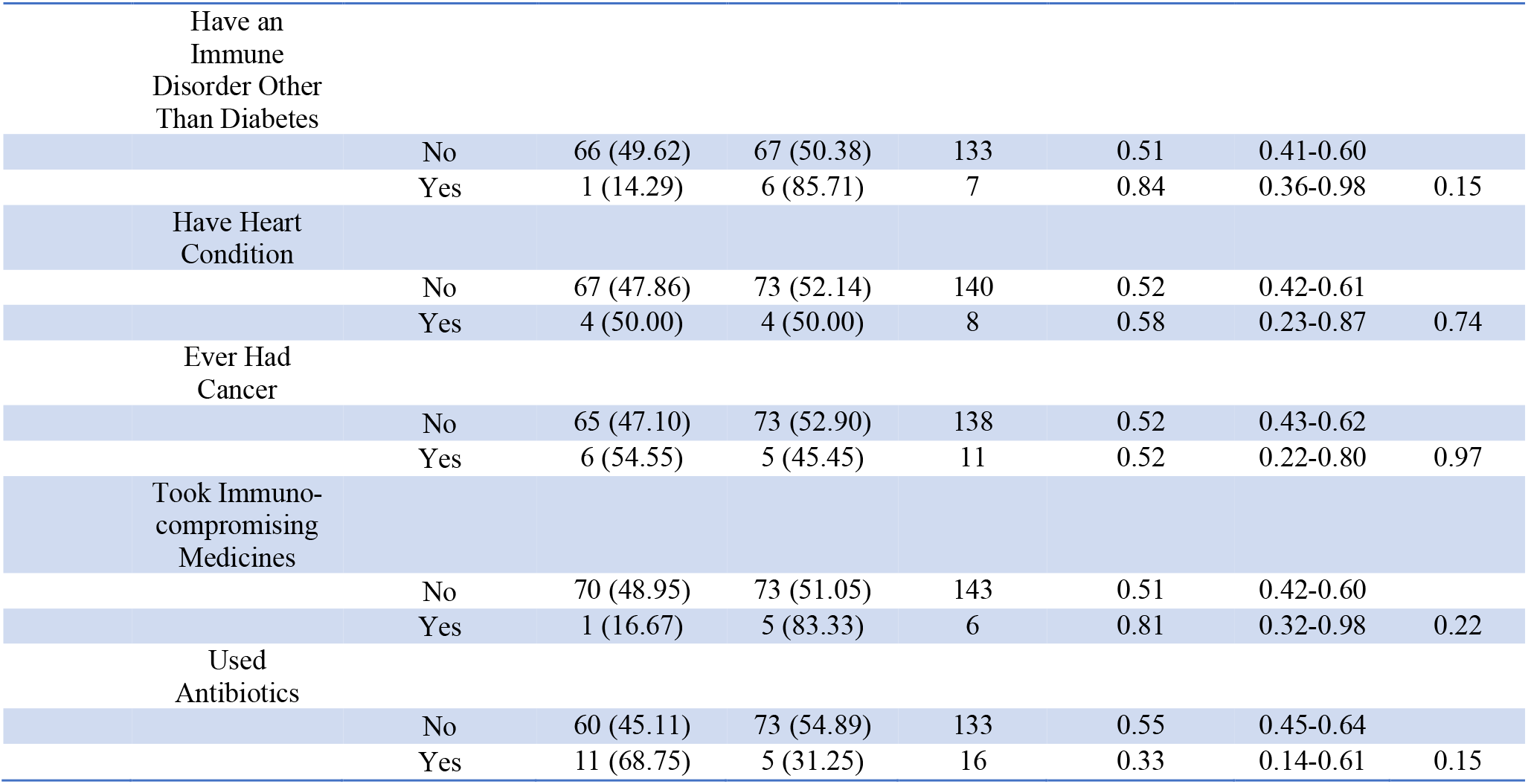
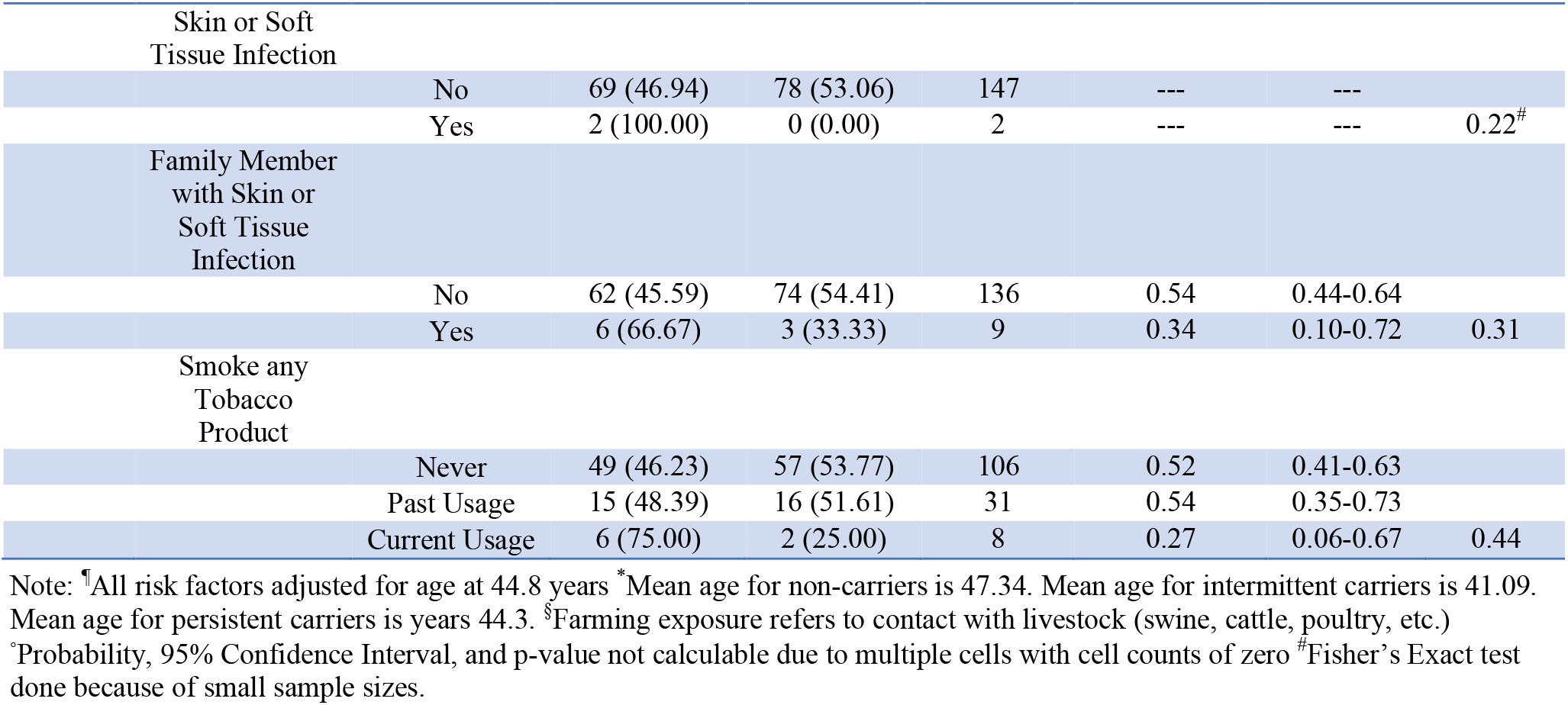
All assessed risk factors for adults

**Table C3:**
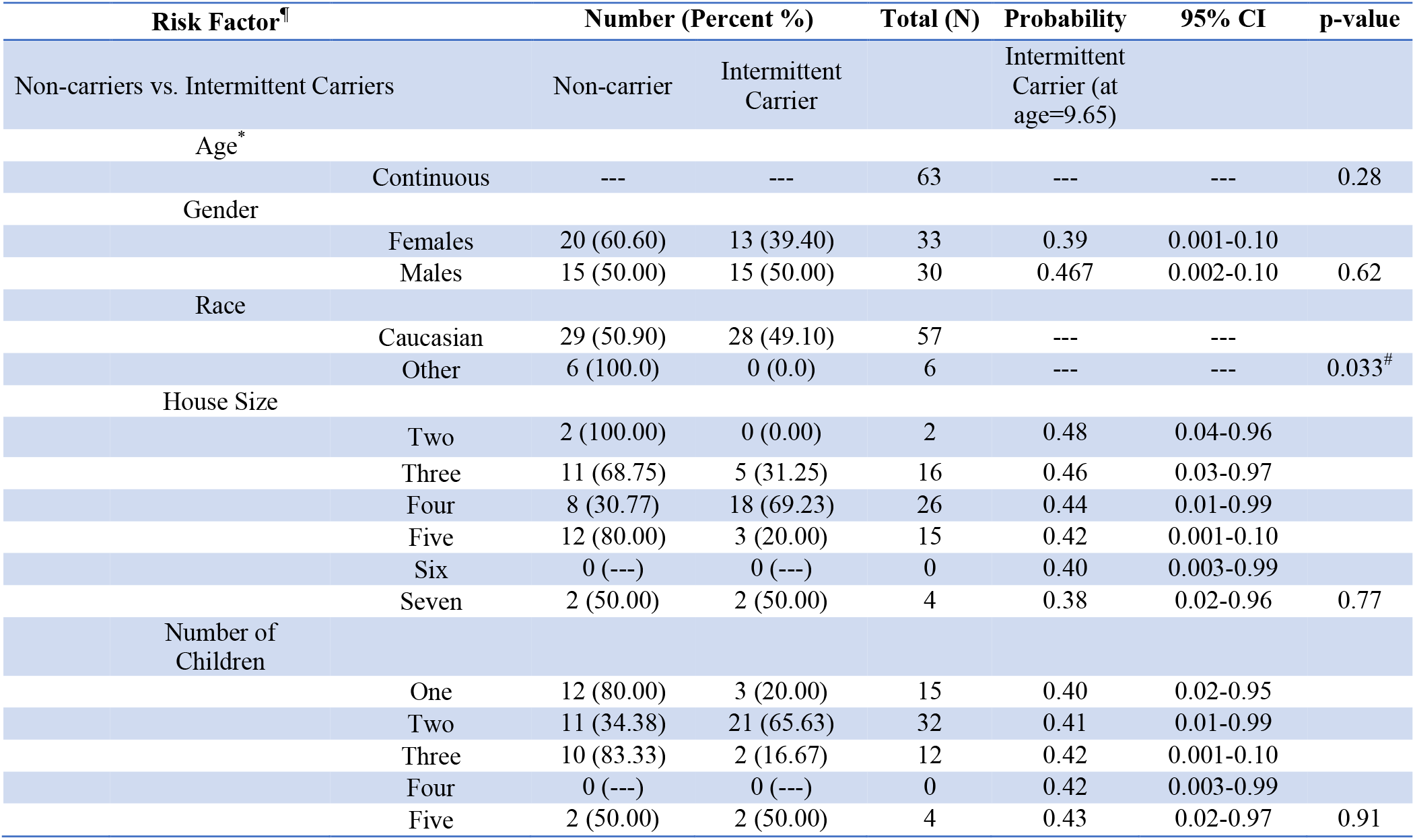
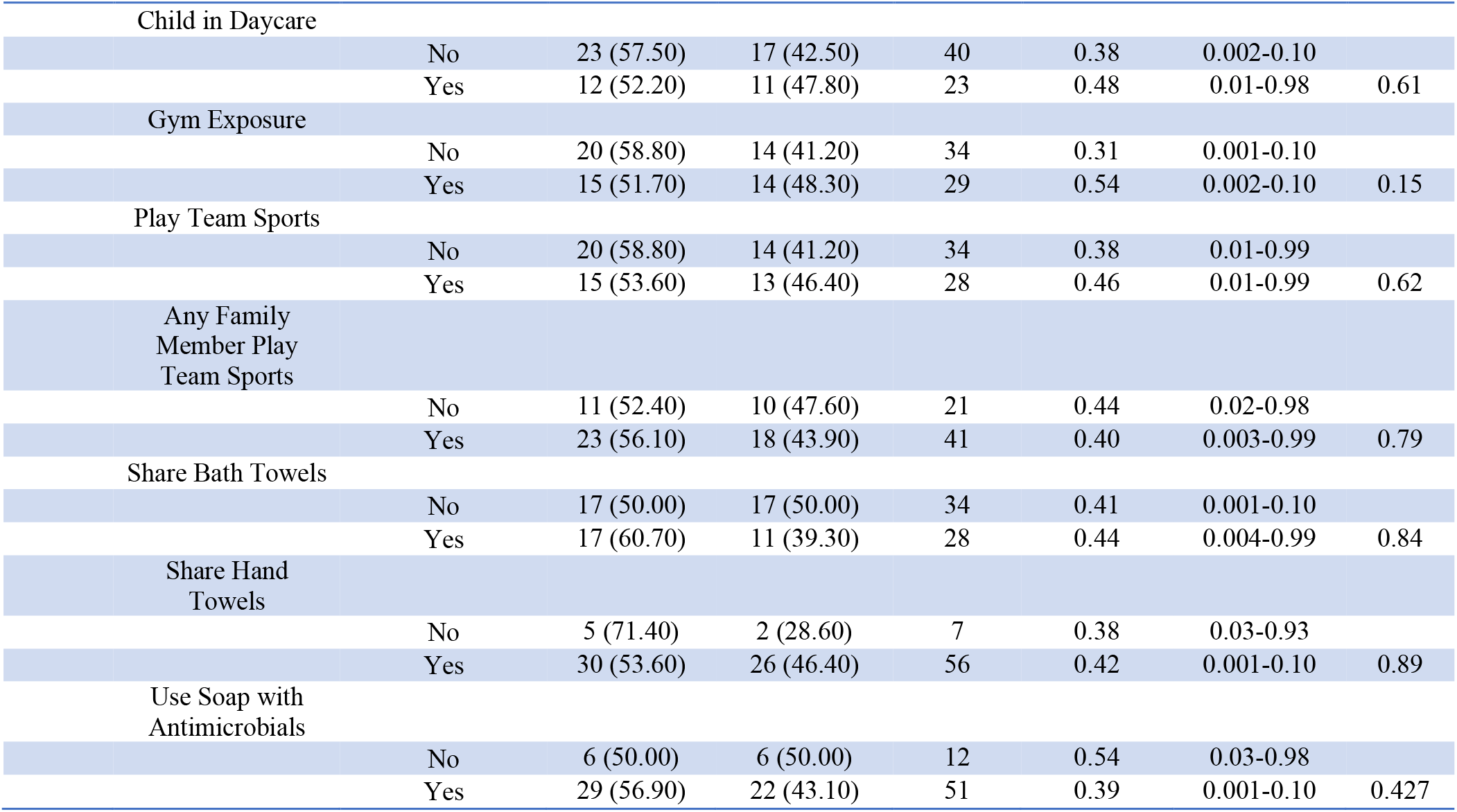
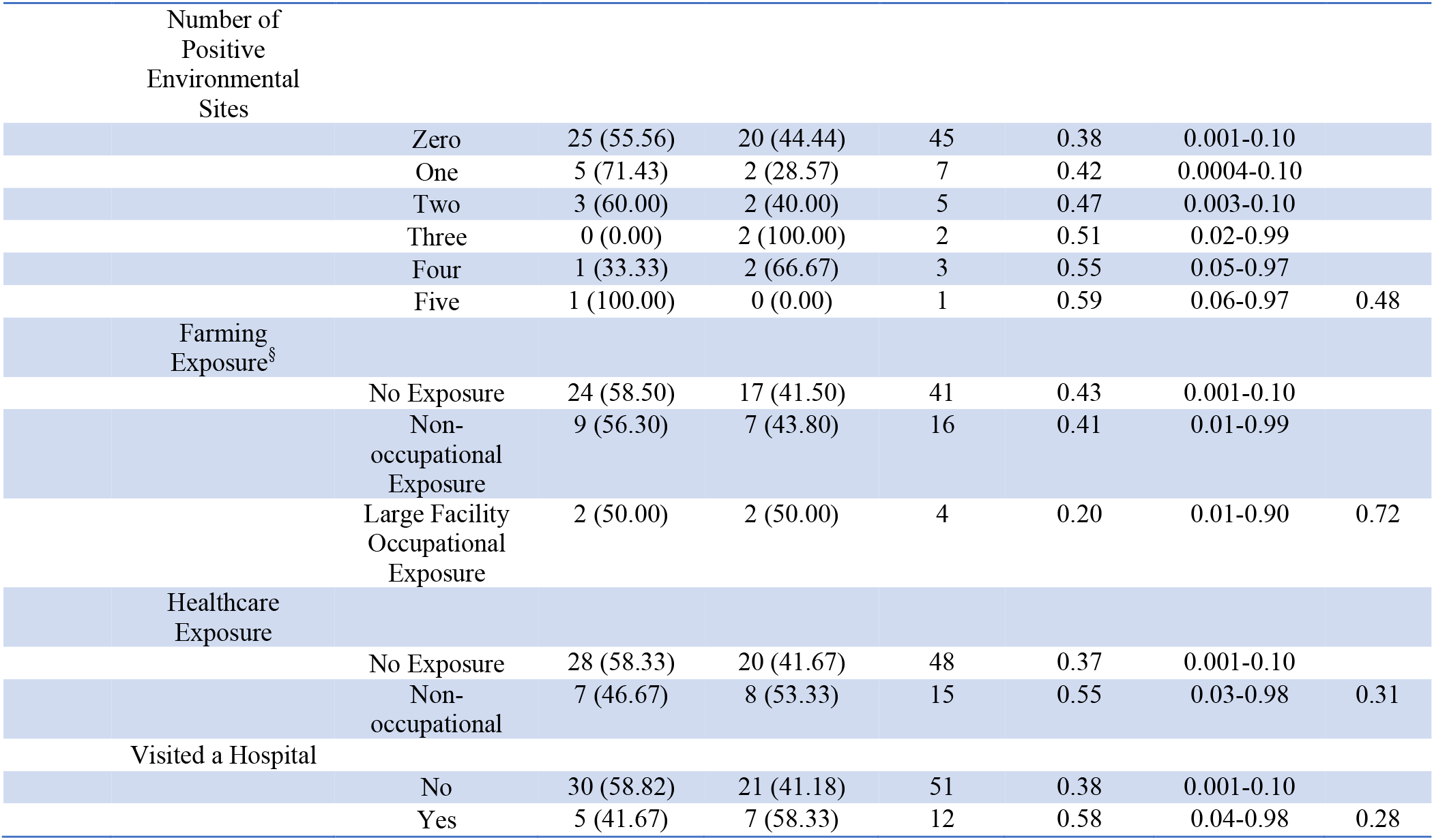
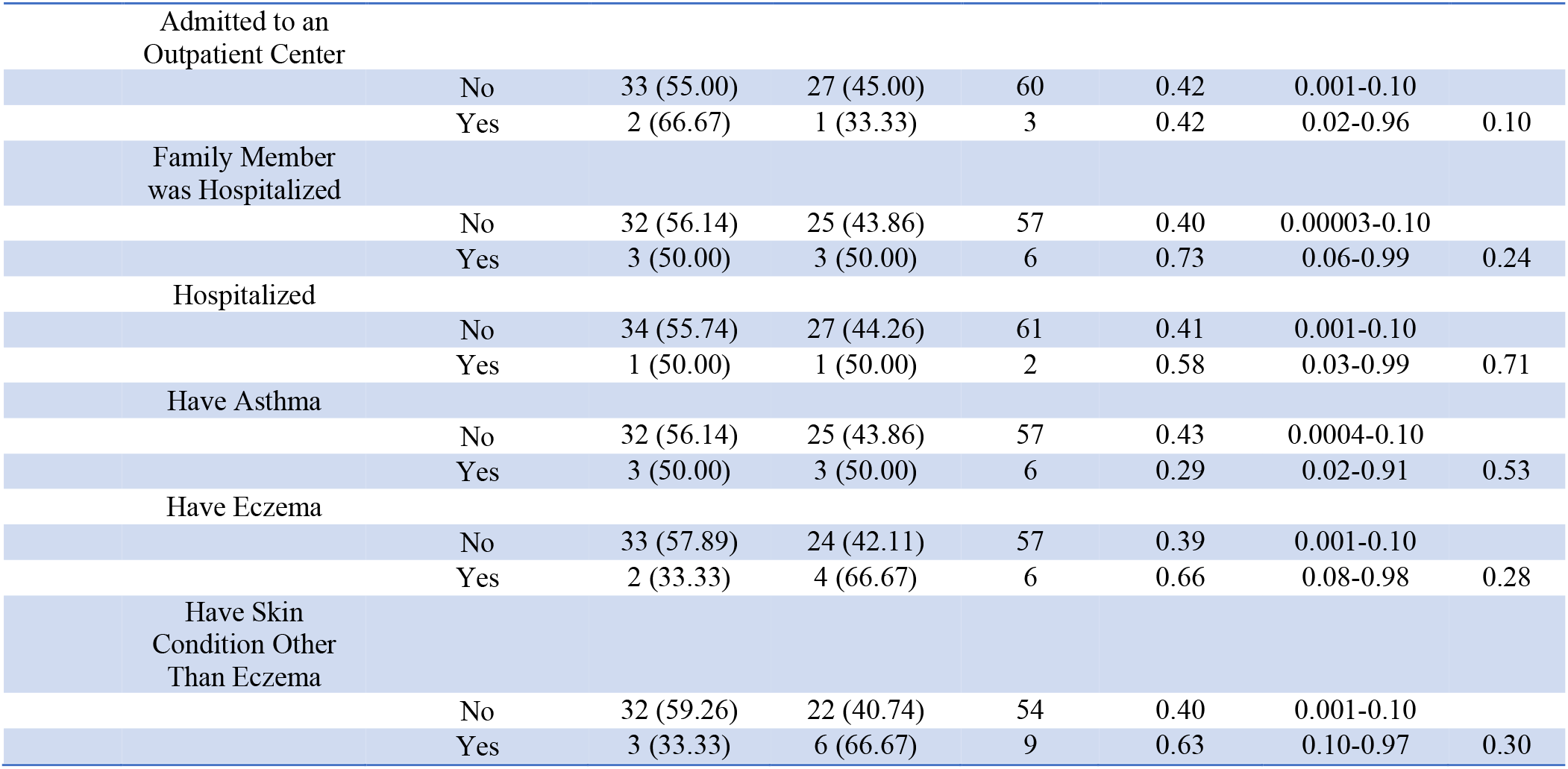
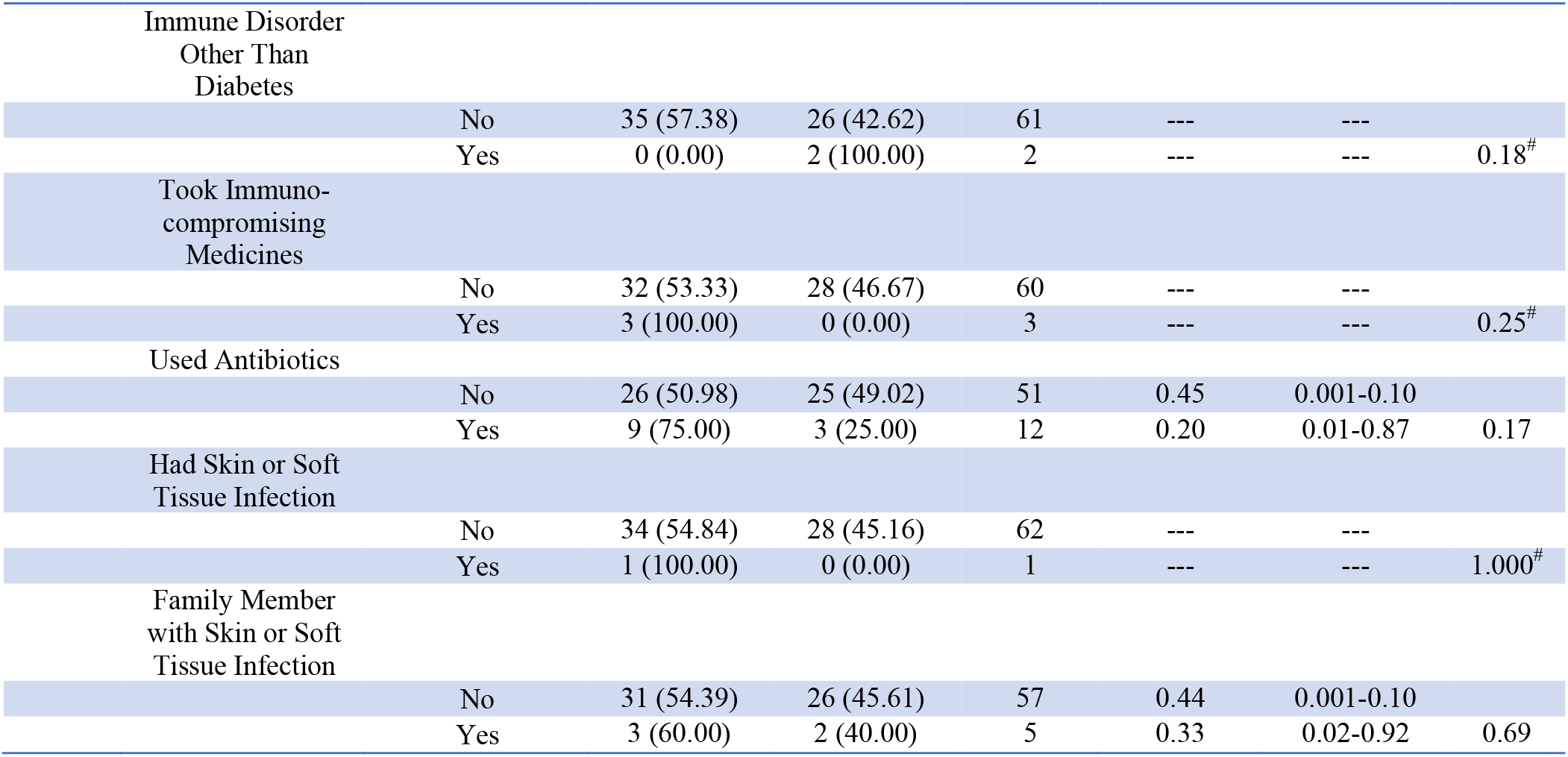
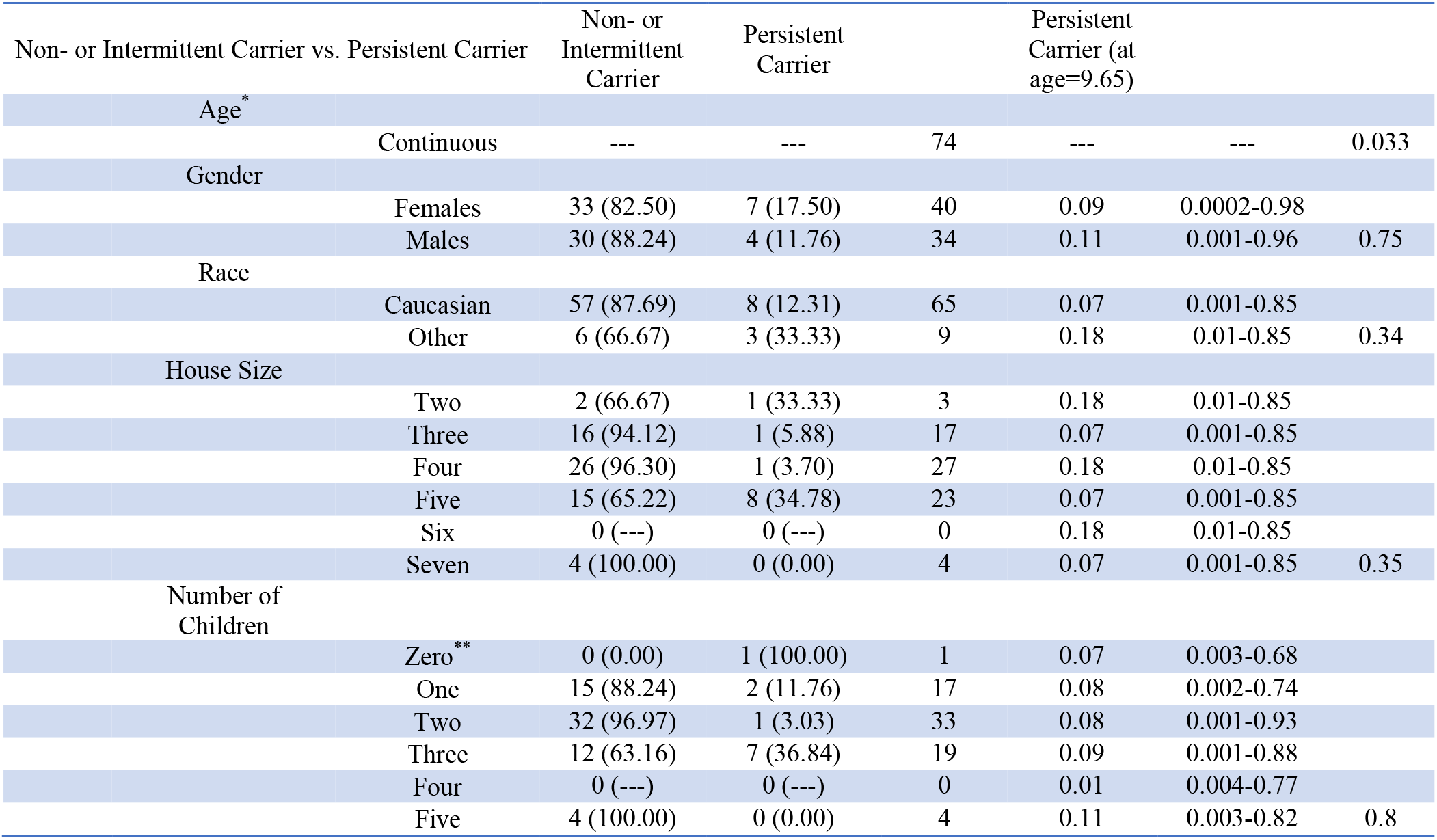
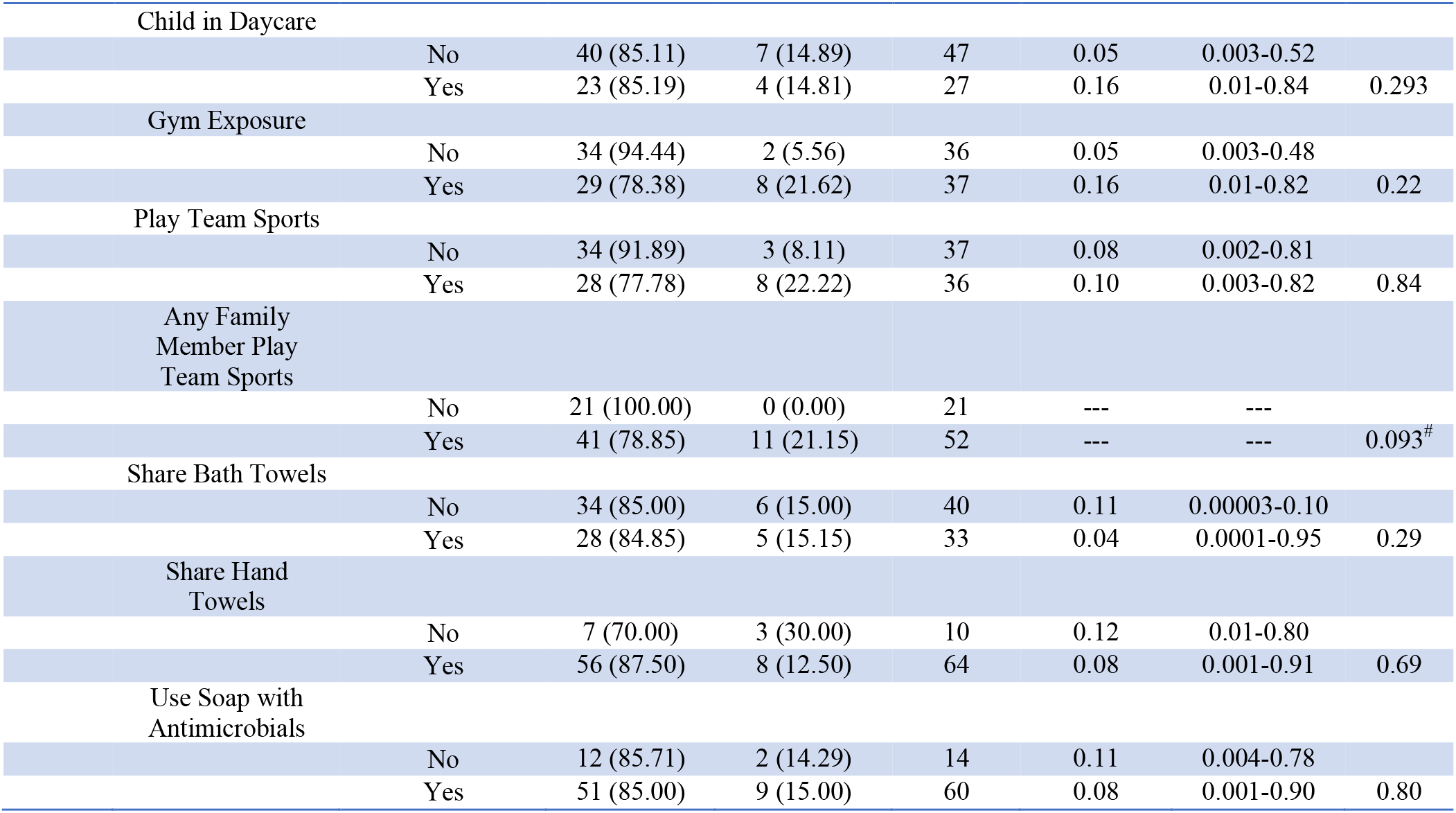
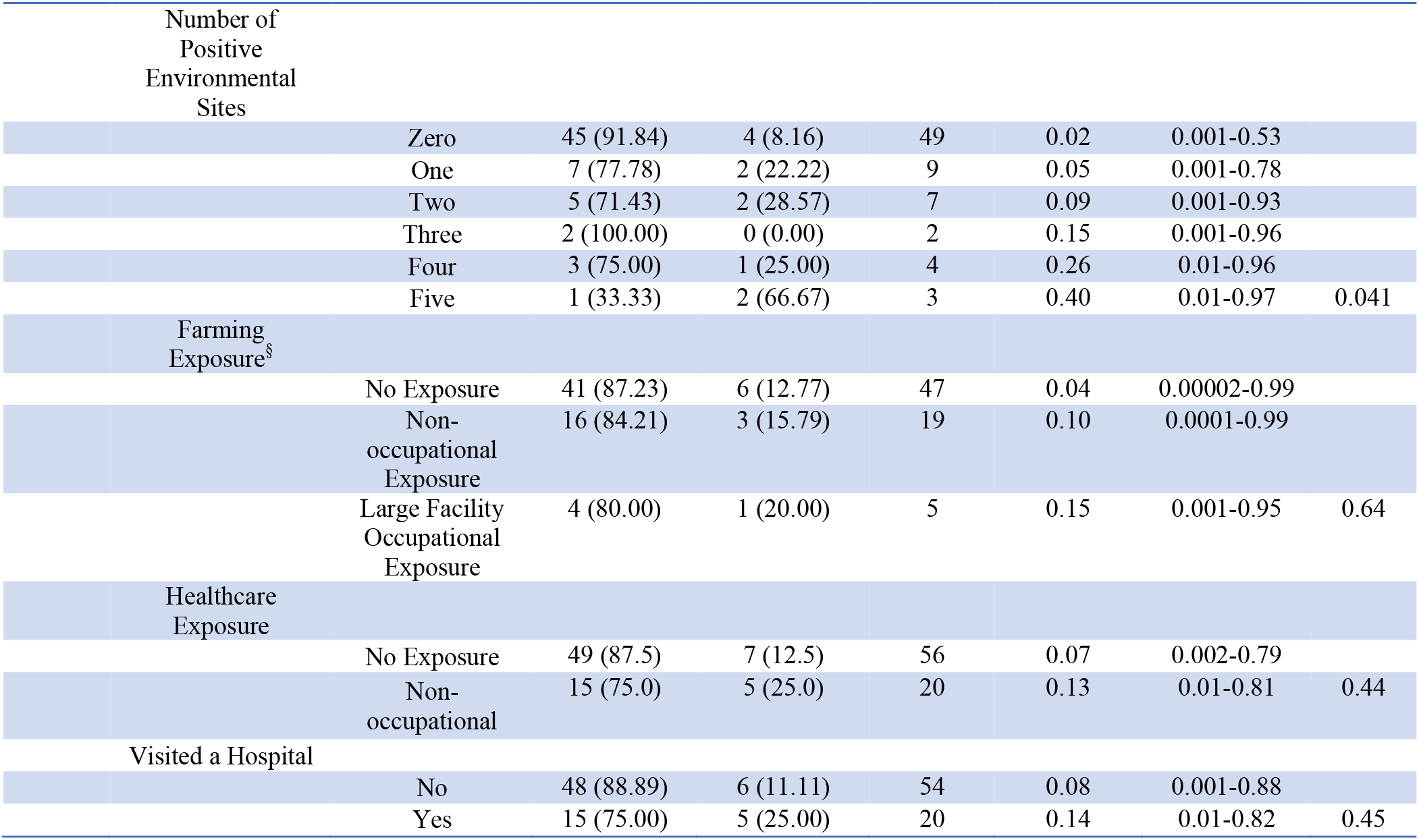
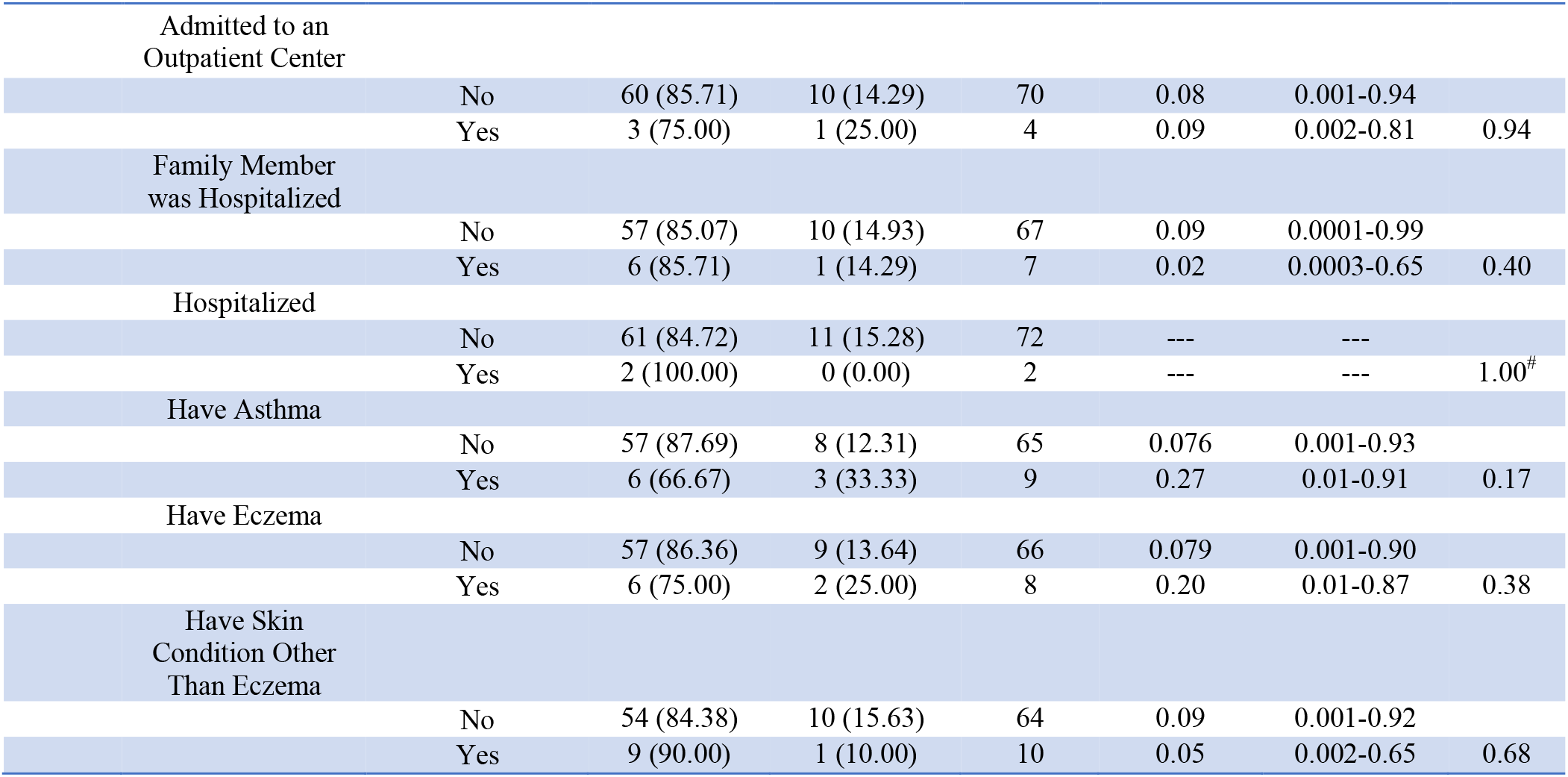
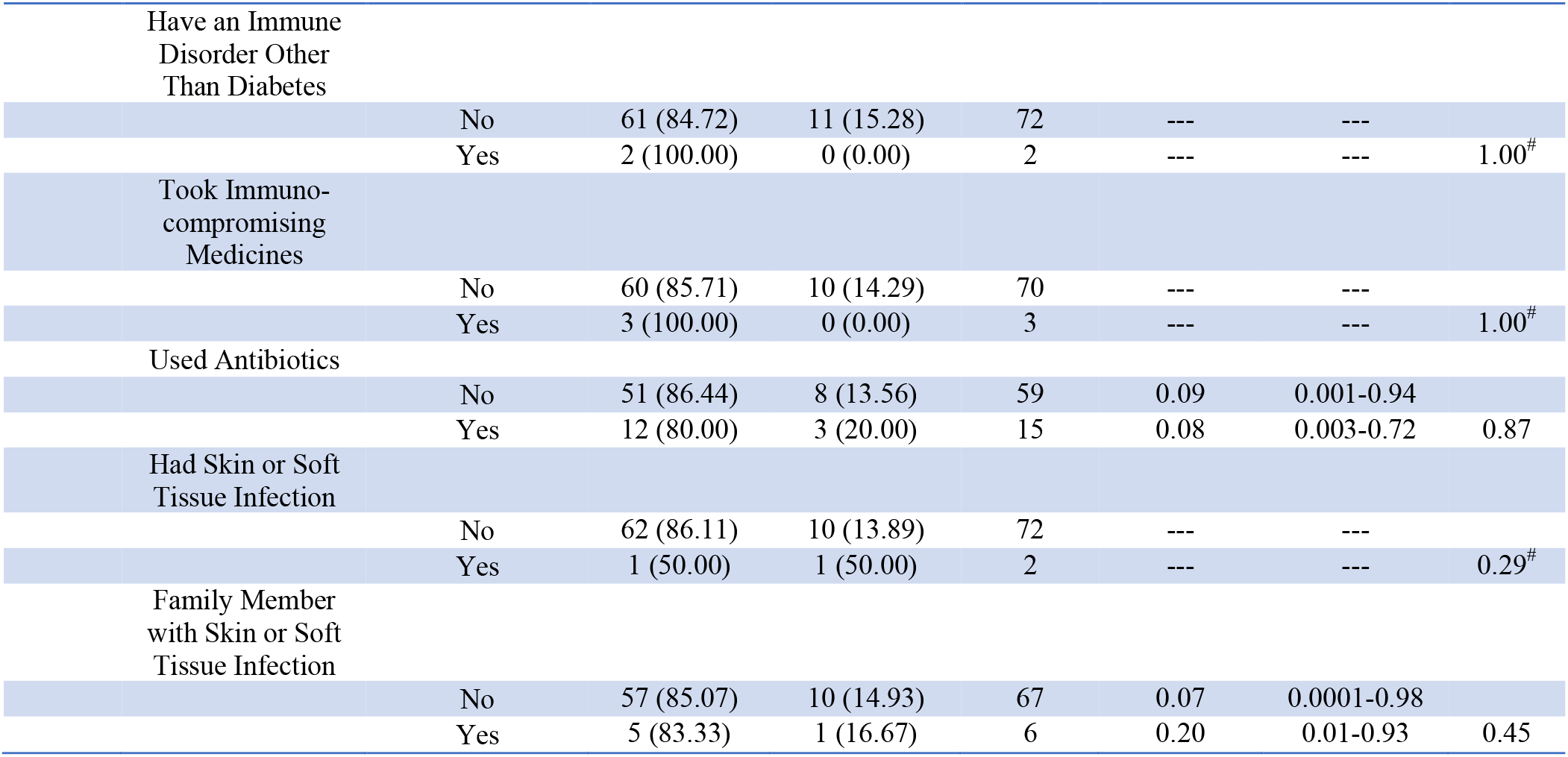
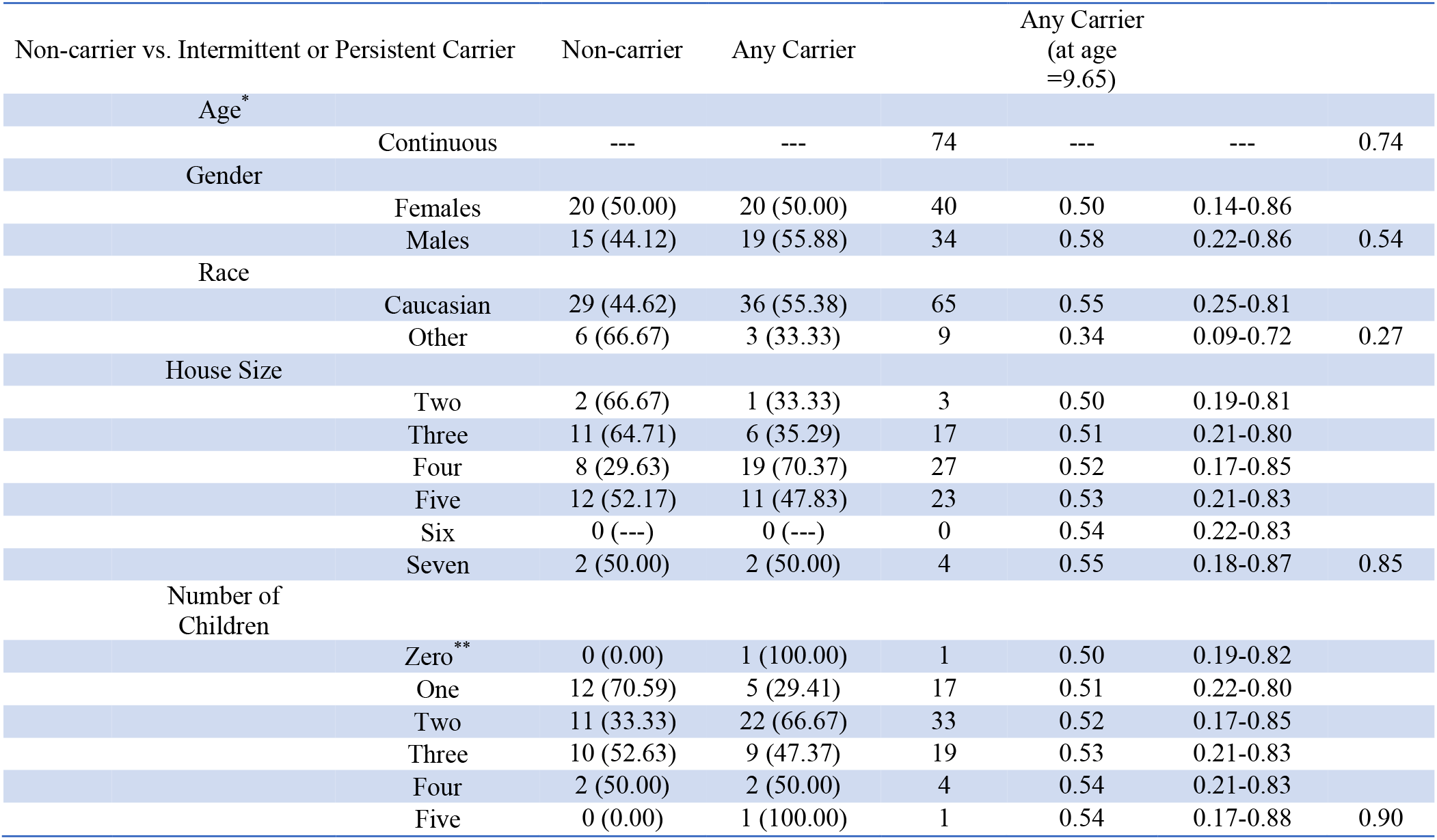
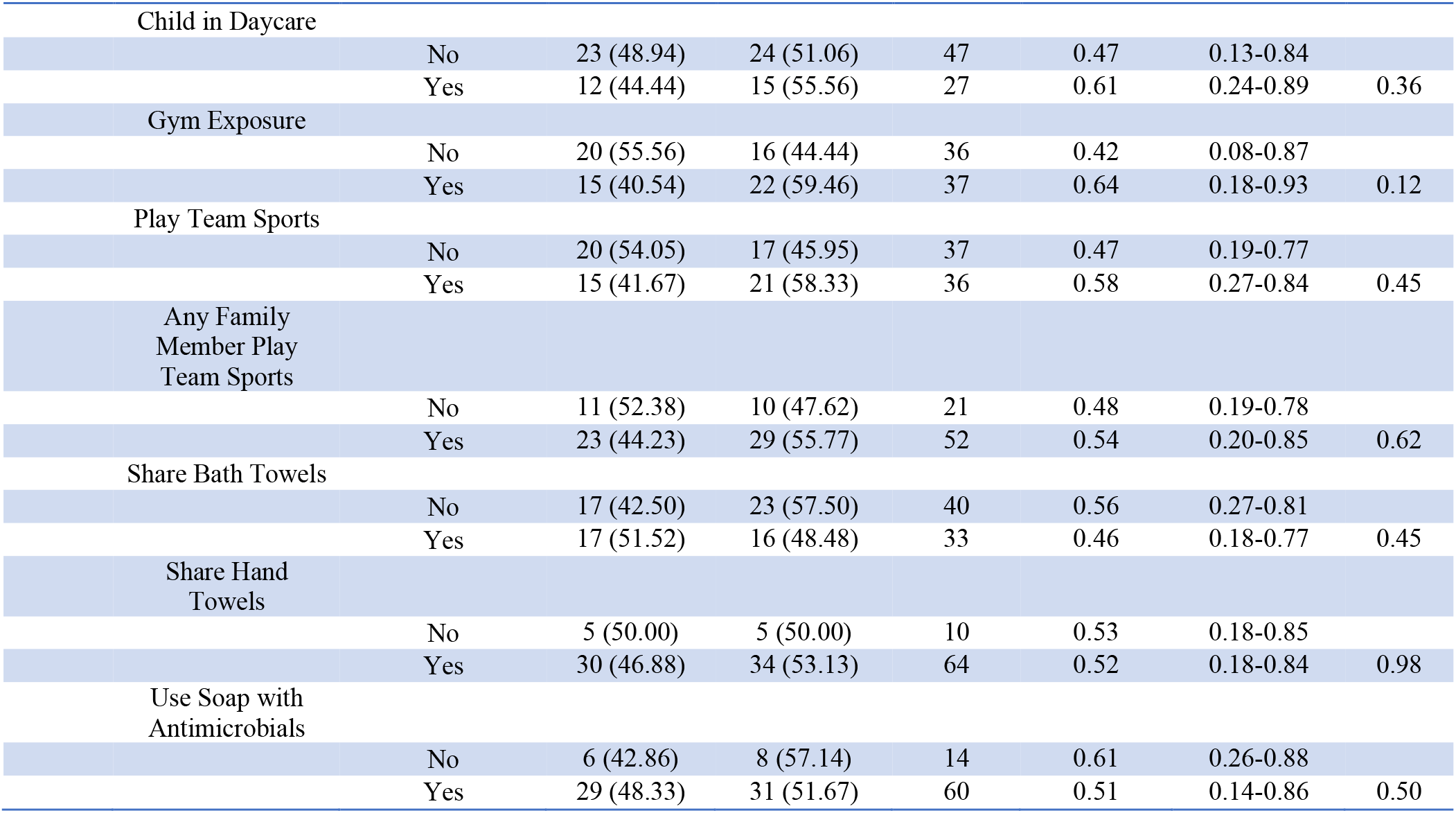
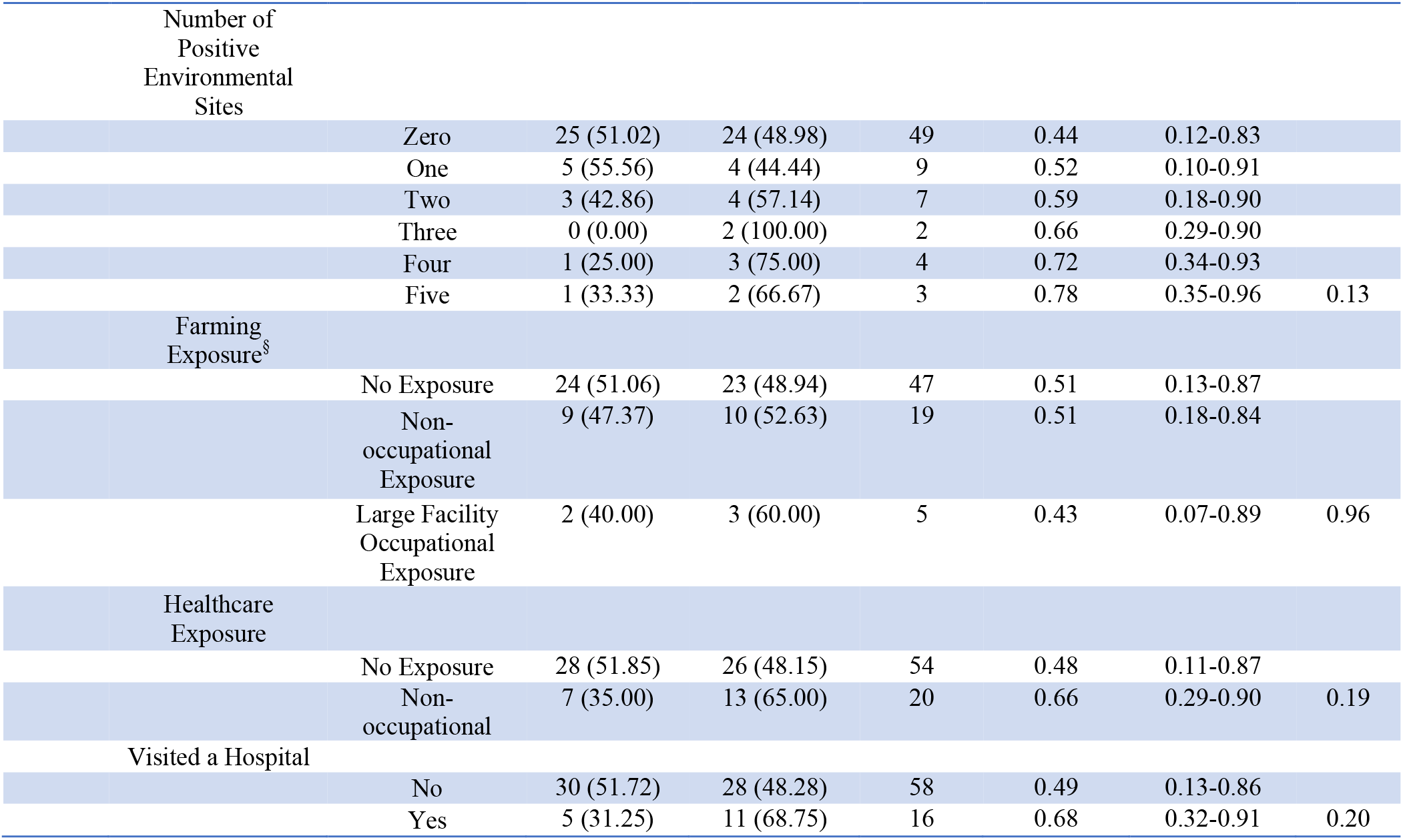
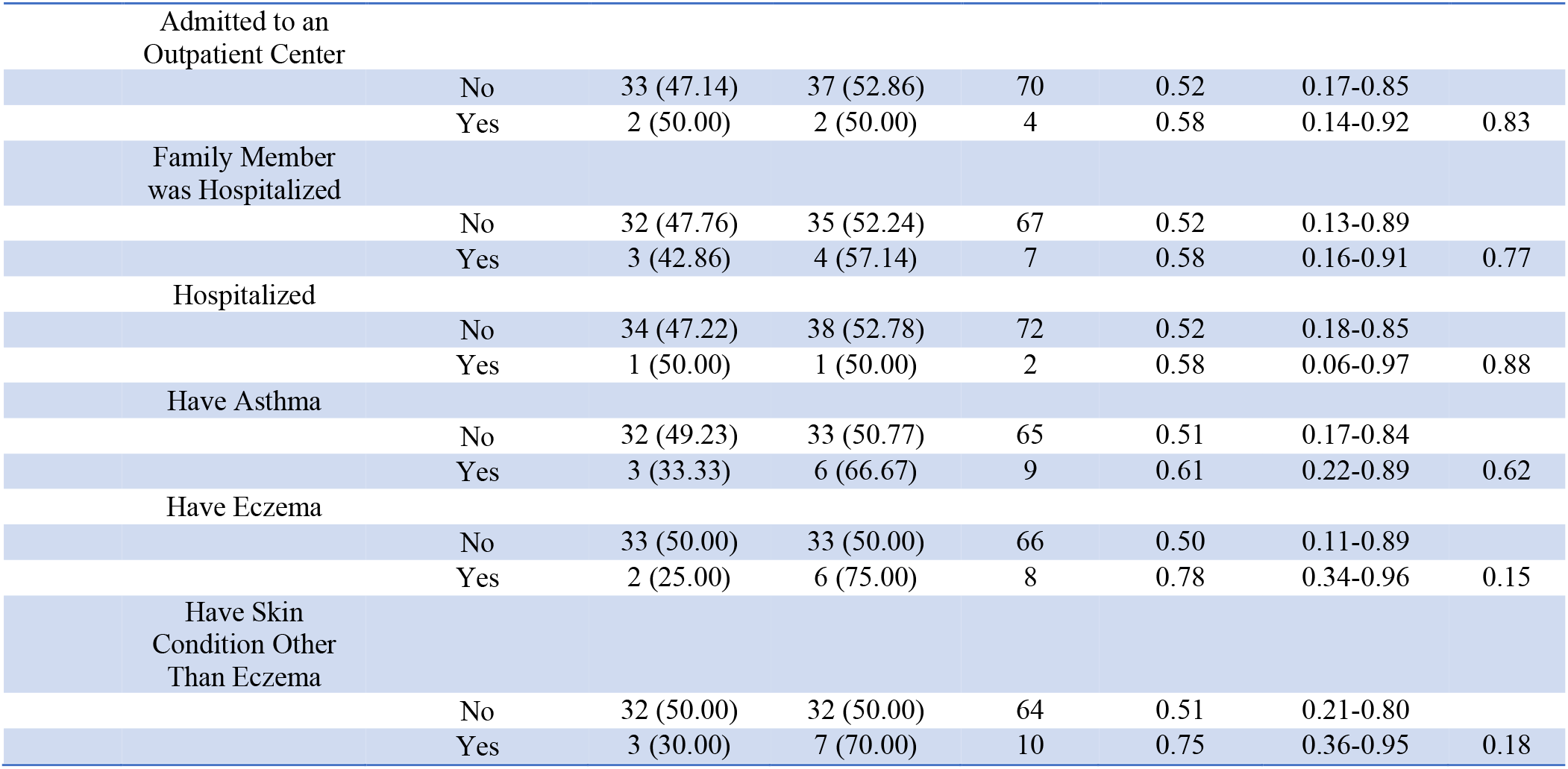
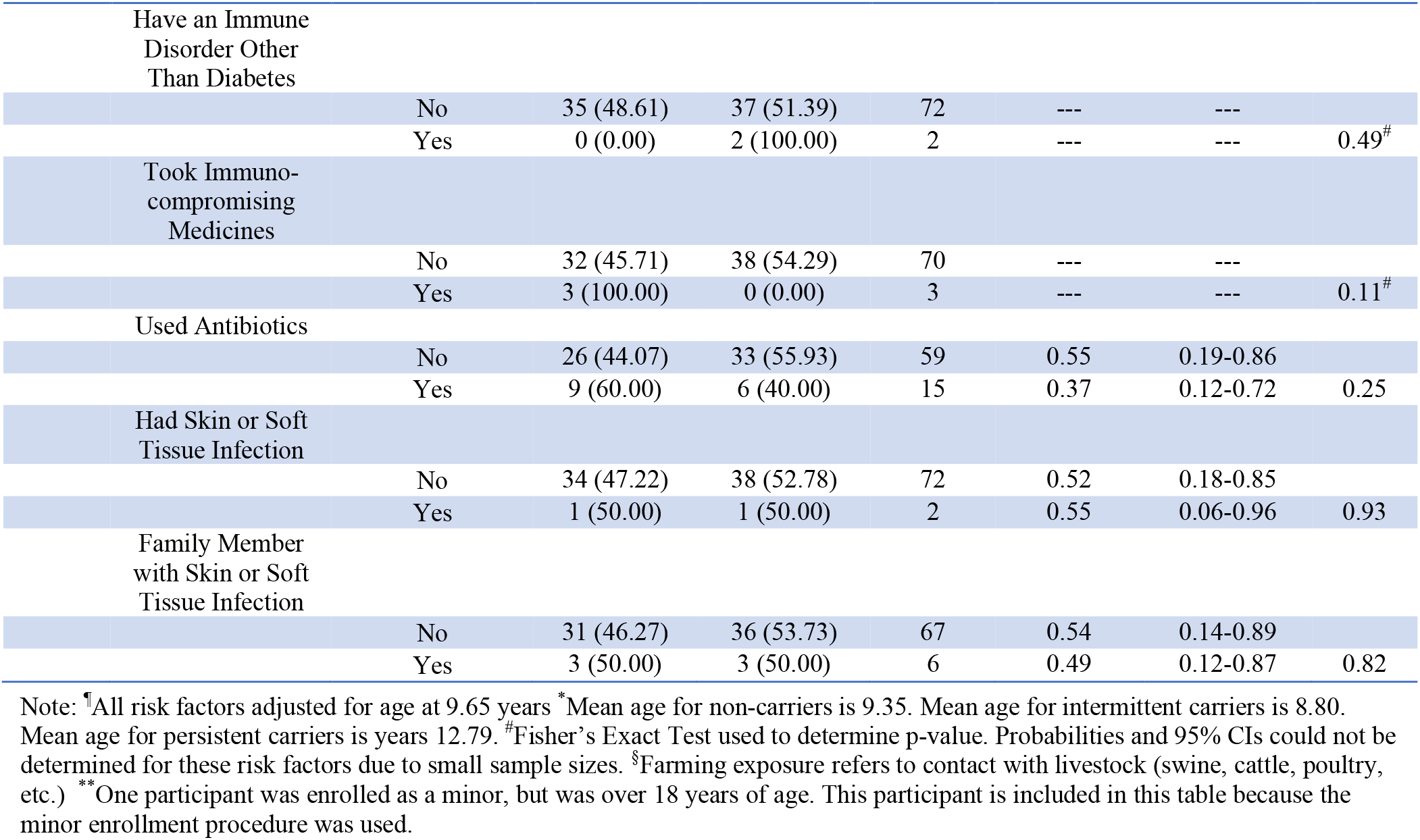
All assessed risk factors for minors

